# Visual Thalamocortical Mechanisms of Waking State Dependent Activity and Alpha Oscillations

**DOI:** 10.1101/2021.04.14.439865

**Authors:** Dennis B. Nestvogel, David A. McCormick

## Abstract

The brain exhibits distinct patterns of recurrent activity closely related to the behavioral state of the animal. The neural mechanisms that underlie state-dependent activity in the awake animal are incompletely understood. Here, we demonstrate that two types of state-dependent activity - rapid arousal/movement related signals and a 3-5 Hz alpha-like rhythm - in the primary visual cortex (V1) of mice strongly correlate with activity in the visual thalamus. Inactivation of V1 does not interrupt arousal/movement related signals in most visual thalamic neurons, but it abolishes the 3-5 Hz oscillation. Silencing of the visual thalamus similarly eradicates the alpha-like rhythm and perturbs arousal/movement-related activation in V1. Finally, we observed that whisker movement or locomotion is not required for rapid increases in cortical activation. Our results indicate that thalamocortical interactions together with cell-intrinsic properties of thalamocortical cells play a crucial role in shaping state-dependent activity in V1 of the awake animal.

**Highlights:**

- Whisker movements correlate with rapid synaptic activation in V1 and visual thalamus
- Silencing of V1 does not abolish movement related activation in most dLGN or LP cells
- Silencing of visual thalamus strongly reduces movement related activation in V1
- Thalamocortical interactions generate state-dependent alpha frequency oscillation
- Visual thalamic cells exhibit LTS firing during alpha oscillation in the awake mouse

## Introduction

Spontaneous, ongoing activity in the waking brain is highly variable and can be classified according to distinct states and rhythms. These brain states are strongly related to animal behavior such as movements and the level of arousal and appear to be one of the strongest determinants of neural activity across the neocortex (Dolensek et al., 2020; McGinley et al., 2015a; Musall et al., 2019; Nath et al., 2019; Niell and Stryker, 2010; Poulet and Petersen, 2008; Reimer et al., 2014; Salkoff et al., 2020; Schneider et al., 2014; Stringer et al., 2019).

One example of a distinct state in the awake brain is the alpha-rhythm (8-12 Hz) first described by Hans Berger in the late 1920s in the human visual cortex (Berger, 1929). Alpha rhythms are particularly strong during states of mental relaxation and eye closure, although a few studies have also reported increases associated with attention and stimulus presentation (Schmid et al., 2012). The cellular and network mechanisms by which alpha oscillations are generated in the visual system are incompletely understood. Studies suggest that the alpha rhythm may primarily originate in the neocortex in layer 2,3 (Halgren et al., 2019), or layer 5 (Silva et al., 1991). Other evidence indicates that visual thalamic nuclei, such as the pulvinar/lateral posterior nucleus (LP) (Saalmann et al., 2012), and/or the dorsolateral geniculate nucleus (dLGN) (Hughes and Crunelli, 2005) are the primary drivers of the visual cortical alpha waves, with high threshold spike bursting proposed as a cell intrinsic mechanism (Crunelli et al., 2018; Hughes and Crunelli, 2005; Hughes et al., 2004; Lorincz et al., 2009). In mice, a state-dependent 3-5 Hz rhythm shares features of the primate alpha rhythm and has been proposed to be an evolutionary precursor (Einstein, 2017; Senzai et al. 2019; Einstein et al., 2017). The neural mechanism of the mouse 3-5 Hz rhythm is currently unknown.

Another example of a distinct brain state in mouse V1 is the strong suppression of delta (0.5-4 Hz), and increase of gamma (30-60 Hz) power in the local field potential (LFP) during high arousal levels (i.e. large pupil diameters) and/or locomotion, which is associated with an increase in the amplitude of visually evoked responses (Niell and Stryker, 2010; Polack et al., 2013; Reimer et al., 2014). On the level of the membrane potential of excitatory V1 neurons, the transition from stillness to locomotion/high arousal is associated with a change from slowly fluctuating synaptic activity to a steady depolarization (Bennett et al., 2013; Polack et al., 2013; Reimer et al., 2014). Several studies have addressed the neural mechanisms underlying this increase in gamma activity and sensory gain in V1 and have suggested the involvement of neuromodulatory pathways (e.g. noradrenergic and cholinergic) (Fu et al., 2014; Lee et al., 2014; Leinweber et al., 2017; Polack et al., 2013), changes in the activity of intracortical inhibitory networks (Fu et al., 2014; Polack et al., 2013; Reimer et al., 2014), and feedback from secondary motor cortex (Leinweber et al., 2017). These different neural pathways work through kinetically different mechanisms (e.g. metabotropic vs. ionotropic receptors) and understanding these kinetics more fully would narrow the possible underlying mechanisms. Indeed, it is not yet fully known if the mechanisms of cortical activation that are associated with movement results from the activation of movement-related structures, or whether this brain activation represents arousal that is concomitant with movement (McGinley et al., 2015b; Petty et al. 2021).

A possible significant contribution of subcortical structures to state-dependent waking V1 network dynamics has recently been suggested based upon the ability of locomotion to alter the activity of early visual brain structures, such as the retina (Liang et al., 2020; Schröder et al., 2020), the dLGN (Erisken et al., 2014), and the superior colliculus (SC) (Ito et al., 2017; Savier et al., 2019; Schröder et al., 2020). Interestingly, studies in the mouse primary somatosensory cortex (S1) have shown that thalamic activity is necessary for the steady depolarization associated with whisker movement (Eggermann et al., 2014; Poulet et al., 2012), but it is not known whether this reflects a conserved mechanism that broadly applies to arousal/movement related brain state changes.

By using intra- and extracellular recording techniques in awake behaving mice together with pharmacological and optogenetic manipulations, we studied the role of thalamocortical interactions in generating the state-dependent alpha-like oscillation and for regulating brain state changes associated with arousal and movements (facial and limb) in mouse V1. Our results indicate that the alpha-like oscillation in mice reflects a thalamocortical rhythm that involves low threshold Ca^2+^ spike (LTS) mediated action potential (AP) bursting in thalamic neurons, depends upon thalamocortical interactions, and predominantly occurs following a reduction in movement and arousal. Furthermore, we found that visual thalamic activity correlates strongly with rapid arousal/movement-related signals in V1 and appears to profoundly shape these signals in V1. Finally, we observed similar rapid state changes during spontaneous periods of increased arousal (pupil dilation) without overt movement, suggesting that a significant component of these rapid transitions in state-dependent activity may operate independently of movement.

## Results

To study the role of thalamic signaling in regulating state-dependent changes in V1, we measured postsynaptic potentials in layer 2,3 neurons through whole-cell recordings in awake head-fixed mice, while monitoring locomotion (wheel movements), whisker movements (whisker pad motion energy), and changes in pupil area. Whisker pad motion energy is a sensitive and informative indicator of the global movement state, with no movement in the whisker pad associated with complete or near complete behavioral quiescence (e.g. Supplemental movies 1,2). Recent studies employing Ca^2+^-imaging and extracellular electrophysiology recordings reported a strong relationship between spontaneous cortical activity and whisker pad movements (Musall et al., 2019; Salkoff et al., 2020; Stringer et al., 2019). The exact timing and kinetics of whisker associated signals in V1 have not been studied and we speculated that addressing this issue would yield important insights into the mechanism that underlies these signals. Tight coupling within the range of tens of milliseconds or less between the onset of whisker movements and the occurrence of postsynaptic signals, as well as fast rise times, would support the hypothesis that whisker movement associated state changes in V1 are mediated via ionotropic signaling. Such fast signaling, in turn, could originate from structures such as the thalamus (Poulet et al., 2012).

To study the timing and kinetics of whisker associated signals in V1, we monitored the face of the mouse at a rate of 8 msec/frame (125 Hz; see Methods). By relating spontaneous behavior to membrane potential changes, we observed several recurring and identifiable states (Fig. 1 and Supplementary Movie 1): 1) behavioral quiescence with small pupil diameter, typically associated with the strong presence of lower frequency (<4 Hz) oscillations in synaptic activity (Fig.1E, slow oscil. shown in yellow); 2) whisker movements with and without locomotion, associated with rapid and sustained depolarization (Fig. 1E,G, sust.depol. shown in red); 3) cessation, or strong reduction, of motor movements, correlated with a 3-5 Hz oscillation (Fig 1E, 3-5 Hz oscil. shown in blue). Closer inspection of these 3-5 Hz oscillations revealed that they were associated with rhythmic barrages of synaptic activity arriving in V1 neurons (Fig. 1F and S1). Locomotion alone, without whisker movement, was not observed. Finally, on rare occasions, we observed pupil dilations without overt body/whisker movements (see below; Fig. S1, Supplementary Movie 2).

**Figure 1.**
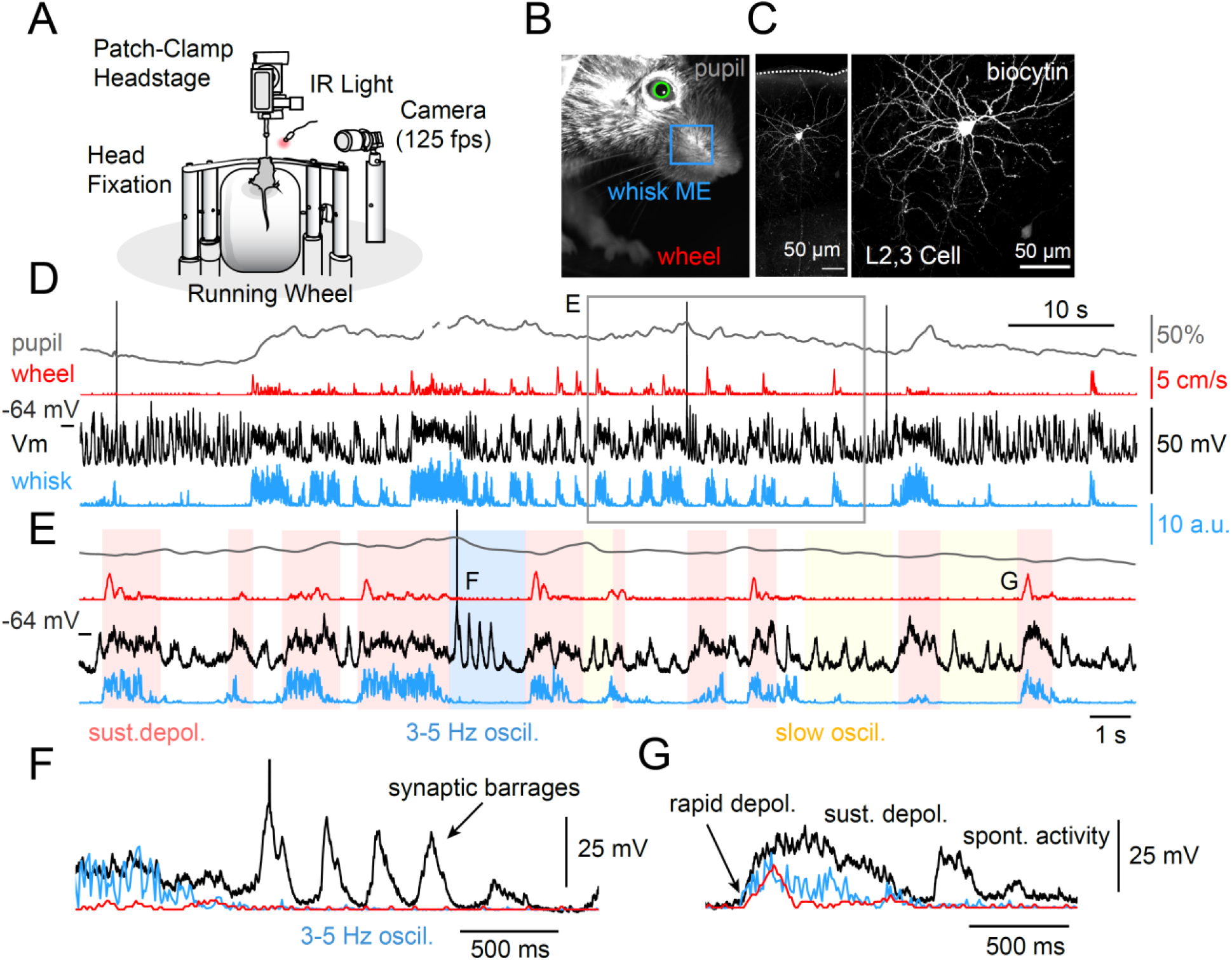
Whole cell recordings from excitatory L2,3 V1 neurons reveal multiple states of synaptic activity. (A) Depiction of experimental setup. The mouse is head-fixed on top of a running cylinder and imaged with infrared light and a camera operating at 125 frames per second. (B) Image of the mouse’s face from the video stream (whisk ME = whisker pad motion energy) (C) Layer 2/3 pyramidal cell recorded and labeled with biocytin. (D) Example whole cell recording from the cell in C. Pupil, whisker pad and wheel movement are also illustrated. Note that sometimes the mouse whisks without walking, but always whisks while walking. Note also that 3-5 Hz oscillations often appear at the end of a movement bout. Pupil diameter follows movement on a slow (seconds) time scale. The membrane potential of this cortical cell depolarizes strongly whenever the animal moves its whisker pad. (E) Expanded time base of a portion of the traces in D illustrating the different phases of activity in this cortical neuron. During the periods labeled in pink, the animal’s whiskers are moving, which is associated with a strong depolarization of the membrane potential of the neuron. During the blue period, the 3-5 Hz oscillation is present, and this often occurs after the cessation of walking. During the unlabeled periods, periodic barrages of synaptic activity recur at lower (<2 Hz) frequencies. (F) Expansion of a 3-5 Hz oscillation illustrating the marked barrages of synaptic activity arriving in this neuron, and the relationship with the cessation of whisker movement. (G) Overlay of the onset of an example bout of whisker movement, walking, and membrane potential illustrating the rapid onset kinetics of the depolarization. Whisker movement precedes walking and appears to be nearly simultaneous with cortical depolarization.

Since the prevalence of slow oscillatory activity during behavioral quiescence has been studied extensively previously (McCormick et al., 2020), we did not examine it further here. Instead, we sought to test the hypothesis that thalamocortical interactions may be responsible for both the rapid depolarization of cortical neurons during arousal and/or movement, as well as the generation of the 3-5 Hz alpha-like rhythm.

### Changes in visual cortical synaptic dynamics associated with whisker movement exhibit rapid kinetics

We recorded from a total of 52 layer 2/3 (L2,3) V1 neurons in 41 mice to study whisker movement/arousal associated membrane potential changes. The physiological features of these cells suggested that they were excitatory neurons (Fig. S3) (McCormick et al., 1985). As in the cell illustrated in Fig.1, we generally observed that the onset of whisking was associated with a rapid depolarization. These rapid depolarizations occurred even in the absence of visible light, which supports the notion that these changes reflect genuine brain state changes as opposed to originating from sensory input at the retina (Fig. S2). In accordance with previous results on locomotion and pupil dilations (Bennett et al., 2013; Polack et al., 2013; Reimer et al., 2014), we found that the membrane potential was less variable during whisker movement (which may also include locomotion) (median 13.67 mV^2^ [19.49-9.85 IQR]); median 12.67 [17.27 - 8.08 IQR]; paired t-test, p = 0.001) and AP firing was significantly increased on average (median 0.36 spks/s [1.08-0.11 IQR].; median 0.54 spks/s [1.59-0.08 IQR]; Wilcoxon signed rank sum test, p = 0.008) (Fig. S3).

Rapid and sustained depolarizations appeared to be more closely coupled, in time, to movement of the whiskers rather than locomotion (which is less frequent than whisker movements), and pupil dilations (which exhibit slow temporal kinetics; Fig. 2). Specifically, the onset of the grand average postsynaptic potential (PSP) response occurred nearly simultaneously, or slightly preceded, the onset of whisker movements (Fig. 2A) with a median half maximum time of 86.09 ms [111.0-30.18 IQR] and a 20%-80% rise time of 158.31 ms [276.93-105.9 IQR] (Fig. 2E,F). We found that PSP responses were more loosely coupled to locomotion onsets with a half maximum time of −49.68 ms [−45.74-(−172.8) IQR] (Wilcoxon rank sum test, p < 0.001) and a 20%-80% rise time of 283.24 ms [416.03-163.23 IQR] (Wilcoxon rank sum test, p = 0.001) (Fig.2 B,E,F). Whisker movement typically preceded the onset of locomotion and we speculate that jitter in the onset of whisk bouts in comparison to locomotion contributed to the slower averaged rise time when PSPs were sorted according to locomotion onsets (Fig. 2B). Both whisker movement and locomotion were associated with pupil dilations and these pupil dilations were less tightly coupled to the visual cortical depolarizations than movements, presumably resulting in part from the slower kinetics of pupillary changes (Fig. 2A,B; Fig. S3).

**Figure 2.**
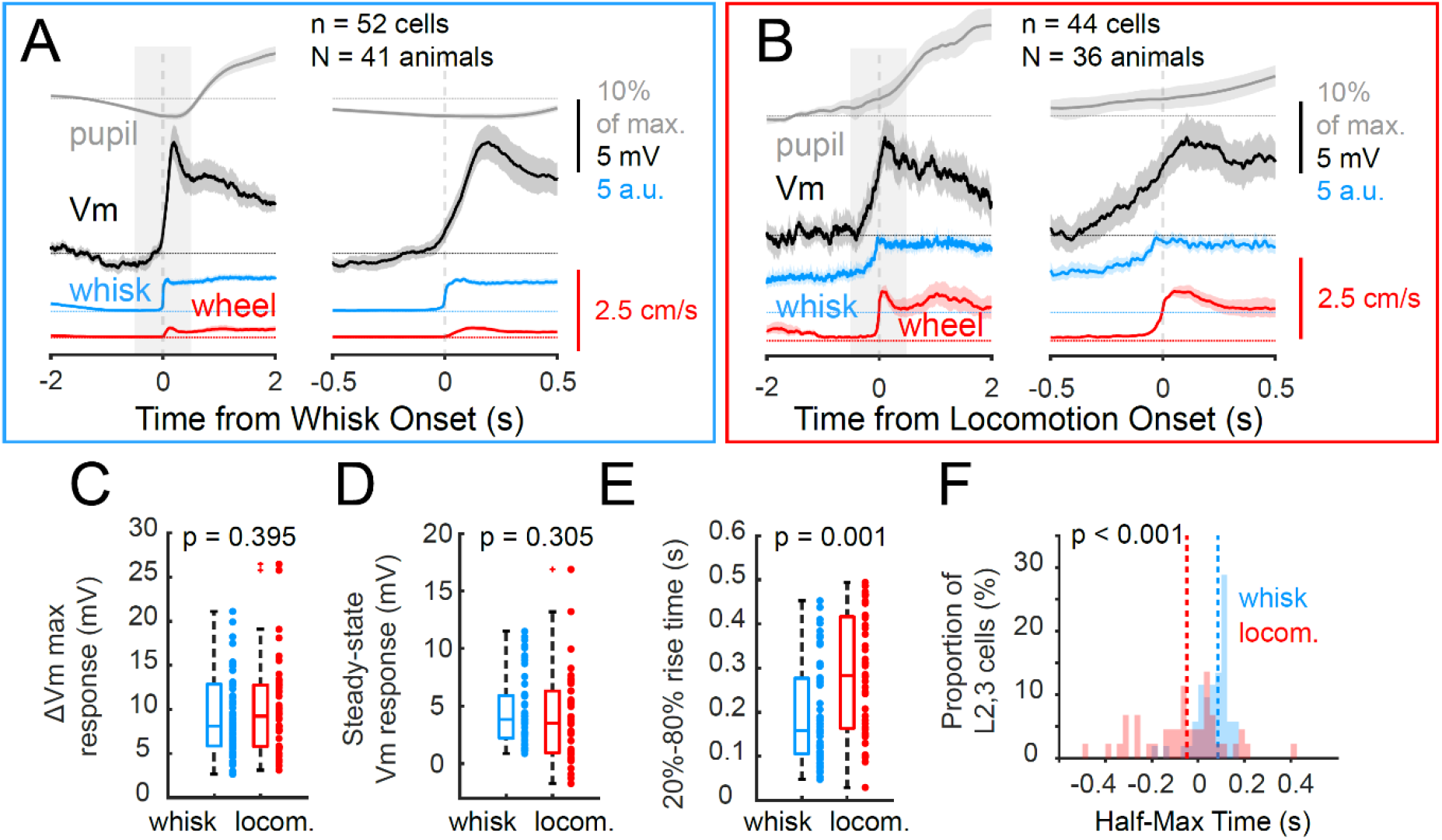
Rapid depolarization of V1 L2,3 neurons is synchronized to the onset of whisker movements. (A) Alignment of recordings to the onset of a whisker movement-bout reveals a sharp synchronized onset of depolarization (two time scales, left and right; n = 52 neurons; N = 41 mice). (B) Alignment of the recording to the onset of locomotion reveals less precise alignment (two time scales illustrated; n = 44 cells in N = 36 mice). (C) Maximum change in membrane potential, (D) steady state depolarization amplitude (between 0.5-2s after movement onset) and (E) 20-80% rise time for a bout of whisker movement or locomotion. (F) Distribution of half maximum time for depolarization for bouts of whisker movement or walking. Negative numbers indicate depolarizations that precede movement. Shaded regions in graphs indicate 95% bootstrapped confidence interval. Boxplots show median, 25th and 75th percentiles and the whiskers indicate the most extreme data points that are not considered outliers. The unpaired t-test (for normally distributed data), or the Wilcoxon rank sum test (non-parametric) were applied to test statistical significance. See also Figures S1,S2 and S3.

On rare occasions, we observed pupil dilations in the apparent absence of whisker movement or locomotion (Fig. S1, supplementary movie 2). We studied this phenomenon in greater detail by visually screening the recordings of 15 randomly selected cells and detected a total of 16 instances in 5 mice (Fig S1). Interestingly, these rare instances of arousal without overt movement were also associated with rapid depolarization of visual cortical neurons, indicating that movement per se is not required for this phenomenon to occur.

What are the neural mechanisms that underlie the rapid state changes in V1? Previous studies have shown that the thalamus plays a crucial role in regulating whisker movement-related depolarization in S1 (Poulet et al., 2012), but it is not known whether the visual thalamus plays an important role for movement related state-changes in V1. Extracellular recordings have revealed a fraction of dLGN neurons to display transiently enhanced AP firing at locomotion onset (Erisken et al., 2014), and for LP visual thalamic neurons to discharge in relation to whisker movements (Petty et al.), but the kinetics of these state changes are not fully known. To address this question, we performed intracellular recordings in excitatory layer 2,3 V1 neurons and combined these with high-density Neuropixels probe extracellular recordings in the two major visual thalamic nuclei (dLGN and LP).

Results from these experiments revealed that whisker movement led to an increase in the average AP firing rate of both dLGN and LP neurons (Fig. 3H), with a similar time course at whisk onset as the depolarizations in V1 (Fig. 3A-G). The half-max time of AP firing in dLGN and LP neurons was similar to that of the synaptic barrage response in V1 L2,3 neurons, with a tendency to occur slightly earlier (median dLGN −37.5 ms [87.5-(−287.5) IQR], median LP −112.5 ms [62.5 -(−412.5) IQR], median V1 106.48 ms [133.345-4.275 IQR]), dLGN vs LP, p1 = 0.062, dLGN vs. V1, p2 = 0.31, LP vs V1, p3 = 0.076, Kruskal Wallis test with posthoc Bonferroni correction) (Fig. 3G). These results indicate that neural activity in V1 and visual thalamus exhibit similar state-dependent changes in activity, with similar onset times for movement-related increases.

**Figure 3.**
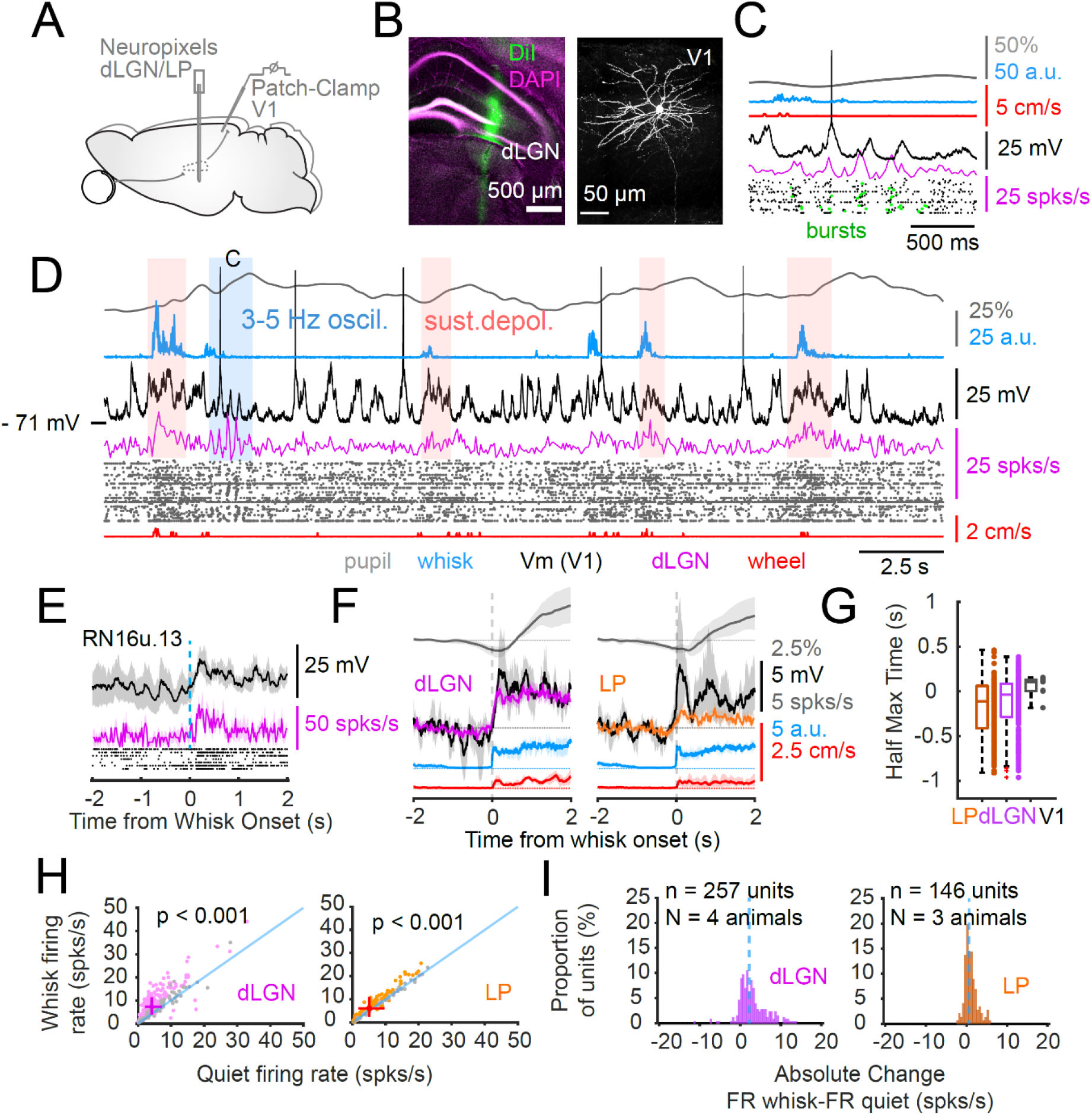
Activity in the visual thalamus is synchronized to the onset of whisker movements and 3-5 Hz rhythmic oscillations. (A) Illustration of recording arrangement with a neuropixels probe inserted into the dLGN and simultaneous whole cell recordings from V1 L2,3 neurons. (B) Illustration of the track of the neuropixels probe (DiI labeling) through the dLGN and a biocytin fill of the simultaneously recorded layer 2/3 pyramidal neuron. (C) Example of the recording, illustrating the occurrence of a 3-5 Hz rhythm that is associated with the arrival of rhythmic PSP barrages in the cortical neuron and synchronized rhythmic discharge of dLGN neurons. Many of these discharges appear as action potential bursts (green dots). (D) Illustration of the recording on a longer time base. Note that when the animal whisks there is a sustained depolarization and increased dLGN activity. (E) Example of a dLGN unit activity (raster and average activity in purple) and V1 neuron membrane potential (black trace; cell in B,C,D) aligned to whisker movement onset. (F) Onset of the grand average of membrane potential change (4 and 3 L2,3 neurons), dLGN (257 units), and LP (146 units) activity aligned to the onset of whisker movement. (G) Comparison of half-max time for dLGN, LP, and V1 neurons. (H) Firing rate of dLGN and LP neurons during periods of whisker movement versus non-whisker movement. (I) Absolute change in activity in dLGN and LP neurons during periods of whisker movement versus non-whisker movement. Shaded regions in graphs indicate 95% bootstrapped confidence interval. Boxplots show median, 25th and 75th percentiles and the whiskers indicate the most extreme data points that are not considered outliers. The crosses in scatter plots show the 25th and 75th percentiles with the center representing the overall medians. The paired t-test (for normally distributed data), or the Wilcoxon signed rank sum test (non-parametric) were applied to test statistical significance in H. Units that were not significantly modulated by whisker movement are shown in grey. The Kruskal Wallis test with posthoc Bonferroni correction was used for data shown in G, which did not reveal any statistical significant difference between the groups.

To determine the membrane potential dynamics of thalamic neurons during rapid state changes, we next performed intracellular recordings from neurons in the LP and dLGN (n = 6 LP cells, n = 2 dLGN cells) (Figs. 4, 5). These recordings, similar to the data from the extracellular high-density neural recordings, exhibited increased AP firing during whisker movement (median quiet 5.26 spks/s [9.45-2.98 IQR]); median whisk 7.97 spks/s [13.56-5.71 IQR], p = 0.004, paired t-test) (Fig. 4G, Fig S4). Whisker movement and locomotion did not significantly alter the variability of the membrane potential (median quiet 7.21 mV^2^ [9.35-6.68 IQR]; median whisk 8.03 [9.91-6.23 IQR]; p = 0.702, paired t-test) (Fig. 4H,I), however, whisk onsets were associated with depolarizations that were reminiscent of the PSP responses described above for L2,3 neurons (Fig. 4E). The depolarizations had a median peak response of 4.59 mV [5.53-2.89 IQR], a median 20%-80% rise time of 273.04 ms [394.98-140.35 IQR] and a median half maximum time of 11.05 ms [60.42-(−102.6)IQR]. (Fig. 4 I-M). These results are an agreement with our findings from extracellular recordings described above and they reveal that the kinetics and the timing of the depolarizations are similarly tightly coupled to whisk onsets in visual thalamic neurons and V1 L2,3 cells. These similarities, in turn, support the possibility that the rapid movement/arousal associated depolarizations in V1 may be regulated/impacted by visual thalamic activity.

**Figure 4.**
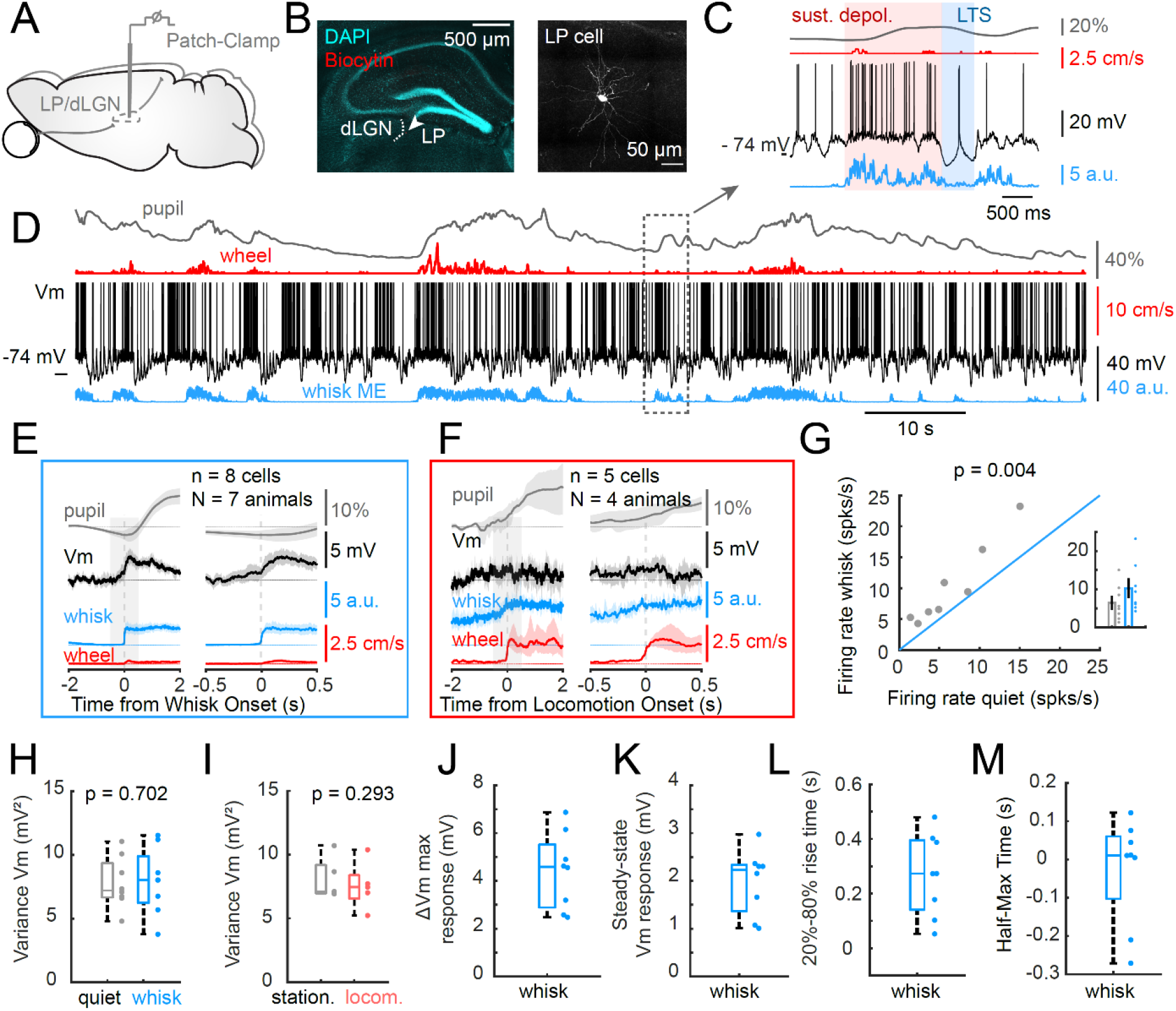
Whole cell recordings from visual thalamic neurons reveal membrane potential dynamics associated with state changes and 3-5 Hz oscillations. (A) Illustration of whole cell recording from visual thalamus (dLGN/LP). (B) Histology illustrating the location of the recorded cell just inside the LP and biocytin recovery of morphology. (C) Detail of the recording shown in D. illustrating a depolarization associated with whisker movement (in red). A short 3-5 Hz oscillation with a low threshold Ca2+ spike (in blue) occurred upon reduction of whisker movement. (D) Longer time base illustrating the activity of the LP neuron shown in B. in relationship to pupil diameter, locomotion and whisker movement. Note the sudden membrane potential hyperpolarizations in thalamic neurons, which were often associated with 3-5 Hz oscillations. Alignment of the membrane potential to onset of whisker movement (E) demonstrates a rapid depolarization, while alignment with the onset of walking (F) does not. (G) The overall AP firing rate is significantly increased in the intracellularly recorded visual thalamic neurons during whisker movement. The membrane potential variance does not significantly change with whisker movement, or locomotion (H,I). Whisker movement-associated maximum changes in membrane potential (J), steady state depolarization (K) and 20-80% rise time (M). L. Distribution of half max rise times for membrane potential changes in visual thalamic neurons (n= 8 neurons, N = 7 mice). Negative values indicate Vm depolarization prior to whisk onset. See also Figure S4A. Shaded regions in graphs indicate 95% bootstrapped confidence interval. Boxplots show median, 25th and 75th percentiles and the whiskers indicate the most extreme data points that are not considered outliers.

**Figure 5.**
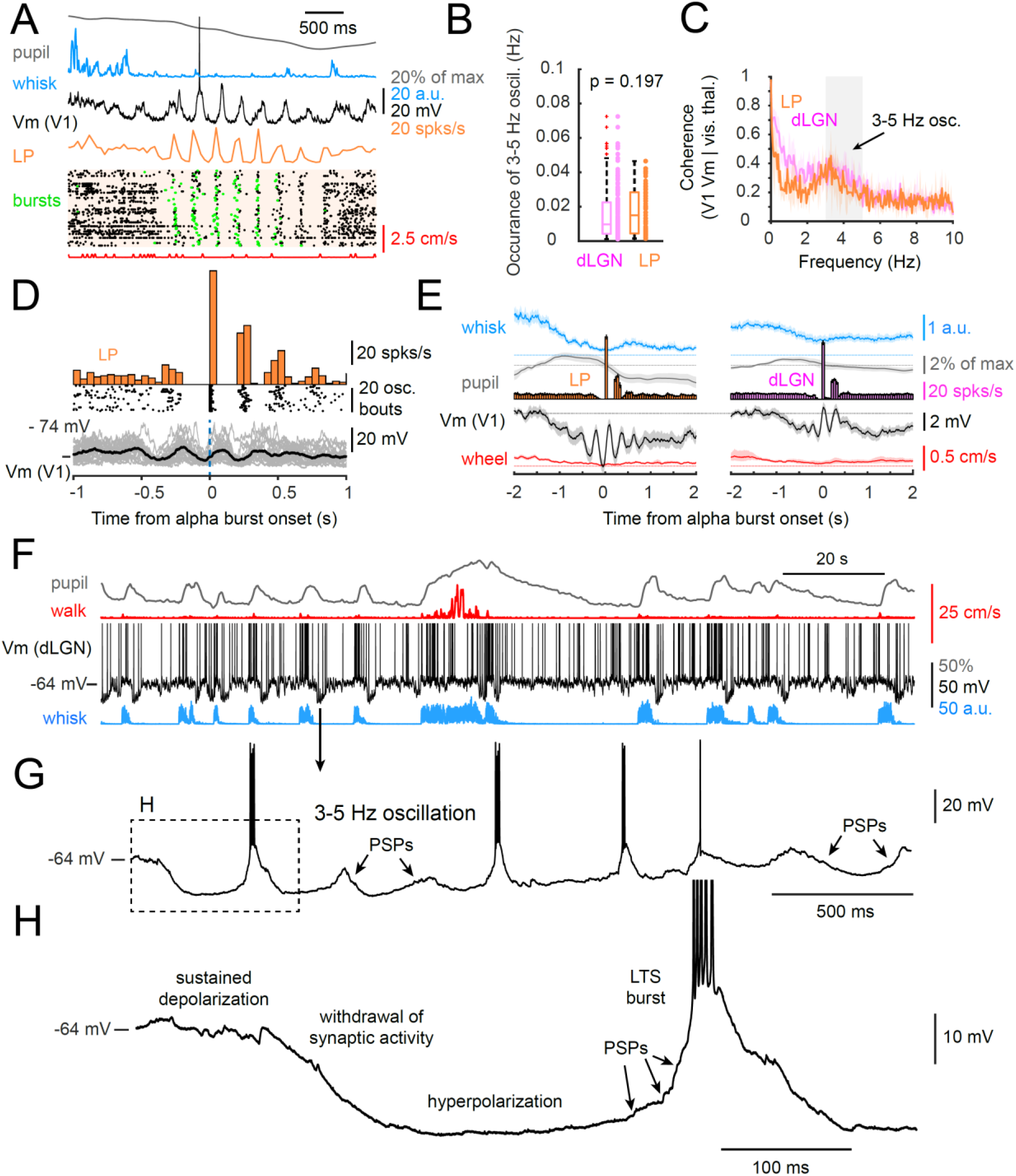
Whole cell and neuropixels extracellular recordings reveal 3-5 Hz oscillations to be highly synchronized in the visual thalamus, and associated with marked oscillations of the membrane potential including low threshold Ca^2+^ spikes. (A) Simultaneous whole cell recording of a V1 neuron with neuropixels recording of unit activity in the LP nucleus during an example 3-5 Hz oscillation. The oscillation is associated with a sudden cessation of activity in LP neurons and a hyperpolarization of cortical neurons, followed by the arrival of rhythmic synaptic barrages and synchronized action potential discharge, including many spike bursts (green dots). (B) The rate of occurrence of the 3-5 Hz oscillation is similar in the dLGN and LP. (C) The membrane potential of V1 neurons and the summed action potential activity of visual thalamic neurons are highly coherent in the 3-5 Hz range. (D) Illustration of the averaged membrane potential of a V1 neuron synchronized to the onset of 3-5 Hz oscillation bouts in an example LP unit. (E) Alignment of grand average of whisker movement, walking, pupil diameter, and L2,3 V1 Vm with onset of 3-5 Hz oscillations in LP and dLGN. (F) Sample recording of a dLGN thalamocortical cell during bouts of whisker movement and locomotion. Note that the membrane potential is always depolarized during whisker movement or walking, but is often depolarized during behavioral quiescence. The 3-5 Hz oscillation appears as a sudden hyperpolarization of the membrane potential - expanded in (G), followed by rhythmic PSP barrages (expanded in H), which often trigger low threshold Ca^2+^ spikes (LTS) (G and H). Shaded regions in graphs indicate 95% bootstrapped confidence interval. Boxplots show median, 25th and 75th percentiles and the whiskers indicate the most extreme data points that are not considered outliers. The Wilcoxon rank sum test was applied for B. See also Figure S4B,C.

### Alpha-like 3-5 Hz rhythm exhibits rhythmic synaptic and intrinsic membrane potential events

Apart from the rapid whisker pad movement associated depolarizations, another common state-dependent pattern of activity in V1 and visual thalamic neurons was the 3-5 Hz alpha-like oscillation (Fig. 1F, 3C, 4, 5, S1). Our intracellular recordings in layer 2,3 neurons of V1 (and L4,5,6; Fig. S1) revealed the 3-5 Hz alpha rhythm to express as an initial hyperpolarization followed by several cycles of depolarization and hyperpolarization, which was also present in neurons from layers 4, 5 and 6 (Fig. S1). Closer inspection of these depolarizing events indicated that they consisted of barrages of synaptic potentials, whereas the hyperpolarizing components appeared as a relative reduction in membrane potential variance, suggesting they represent periods of reduced synaptic activity (Fig. 1F, 3C, 5A, S1). Simultaneous neuropixels probe recording of multiple neuron activity in either the dLGN or LP with simultaneous whole cell recordings in L2,3 V1 neurons revealed prominent synchronized AP activity in the thalamus that coincided with the barrages of synaptic potentials arriving in visual cortical neurons (Fig. 3D, 5A). Examining the spectral coherence between averaged/summed visual thalamic activity and membrane potential oscillations in V1 revealed a prominent peak at approximately 3-5 Hz (Fig. 5C) indicating a strong relationship between these thalamic nuclei and V1 neuronal membrane potential within this frequency range. We further found that the 3-5 Hz rhythm was associated with frequent AP burst firing in dLGN and LP units, characterized by 2-5 APs with intraburst frequencies of 250-350 Hz, and preceded by AP quiescence of greater than 100 msec, which indicates that they are initiated by intrinsic low threshold Ca^2+^ spikes (McCormick and Bal, 1997). We detected at least 3 bouts of such AP bursts at an inter-burst frequency of 3-5 Hz during our ~30 min recording sessions in 176 out of 257 dLGN units (~69%), and in 91 out of 146 LP units (~62%) at median rates of 0.01 Hz [0.023-0.004 IQR] and 0.015 Hz [0.028-0.004 IQR] (Fig. 5B). Sorting the membrane potential of V1 neurons to the onset of these bursts revealed that the oscillation coincided in the visual thalamus with rhythmic depolarisations in L2,3 V1 neurons (Fig. 5D,E).

Averaged whisker movement activity and wheel movements exhibited a decrease preceding the onset of the 3-5 Hz oscillations (Fig. 5E), indicating that this rhythm preferentially occurs after bouts of enhanced movement, during which the pupil constricted (see also (Einstein et al., 2017)).

We detected the 3-5 Hz oscillations in 7 out of our 8 intracellularly recorded visual thalamic neurons. At the onset of these oscillations, the membrane potential of dLGN/LP neurons showed a sudden and pronounced (median, −78 mV [−74.75-(−81.5) IQR] hyperpolarization, which was associated with a decrease in membrane potential variance, suggesting that it is mediated by the sudden withdrawal of depolarizing synaptic barrages (Fig. 5F-H and S4). Following this onset-hyperpolarization, the membrane potential slowly, and then suddenly, depolarized, often initiating an LTS spike crowned by 1-4 APs (Fig. 5G,H). Closer examination of this sudden depolarization revealed it to be mediated by the arrival of PSP barrages, which clearly included EPSPs, based upon their fast rise and slower fall in membrane potential (Fig. 5G,H). The 3-5 Hz oscillation in visual thalamic neurons appeared as several (2-7) cycles of hyperpolarization-depolarization. Often, the depolarizations failed to initiate either a low threshold Ca^2+^ spike or action potentials. In these cases, the barrage of PSPs arriving in thalamic neurons during this oscillation were clear (Fig. 5 G,H and S4). Interestingly, in recordings from LP neurons, these PSPs often contained EPSP-like events that were of larger amplitude (2-10 mV), suggesting they may arise from activity in layer 5 corticothalamic neurons (Sherman and Murray Sherman, 2016). Gradually, over the course of the oscillation, the hyperpolarized membrane potential lessened, and the oscillation stopped (Figs. 4,5). Between bouts of 3-5 Hz oscillation, the membrane potential of thalamic neurons often exhibited a relatively sustained depolarization, often in the absence of overt movement (Fig. 4C,D; Fig. 5F).

Examining the timing of visual thalamic activity (recorded with neuropixels probes) with synaptic activity in visual cortical neurons (whole cell recordings) revealed that the initial silent period of thalamic activity during the onset of the 3-5 Hz oscillation is associated with a sudden hyperpolarization and drop in AP firing in cortical neurons and that the oscillations are synchronized (Fig. 5A; n=7 simultaneous recordings). Since we did not align the visual field locations of the simultaneous thalamic/cortical recordings, we did not attempt to compare precise onset times of the 3-5 Hz rhythms between these structures.

### Optogenetic silencing of visual cortex abolishes 3-5 Hz rhythm

Based upon our recordings, we hypothesized that the arousal/movement-related changes in thalamic and cortical activity, along with the 3-5 Hz rhythms, may represent thalamocortical interactions. To test this hypothesis, we employed an optogenetic approach whereby activity within V1 was suppressed by stimulating channelrhodopsin expressing PV positive interneurons (e.g. (Speed et al., 2019)).

Optogenetic activation of PV interneurons in V1 suppressed AP activity throughout all cortical layers under the center of light delivery (data not shown). Although the stimulation was focused on V1 through a relatively small craniotomy (~300 um), we cannot rule out the possibility that other higher visual cortical regions were also affected. For this reason, we will refer to this manipulation as silencing of the visual cortex, rather than of V1 in specific. Upon suppression of activity in visual cortex, we observed that the median activity in dLGN units was suppressed by −46.55% [−1.22-(−6.94) IQR], while LP units showed an even larger median reduction in their activity by −83.26% [−5.33-(−9.68) IQR] (Figs. 6,7 A-C). These results are consistent with the strong dependence of dLGN neurons upon retinal input, while LP cells appear to be mainly driven by visual cortical activity (Sherman and Murray Sherman, 2016). During the optogenetic silencing of the visual cortex, 3-5 Hz oscillations in the dLGN and LP were abolished (Fig. S5). In contrast, the optogenetic silencing of the visual cortex did not have a uniform effect on whisker movement associated signals in dLGN and LP units (Fig. 6,7D-I). About 25% of the units in the dLGN and ~10% of the units in the LP showed a significant increase in whisker movement related signals during visual cortical silencing (Fig. 6,7F,G). In ~12% of the dLGN units and in ~28% of LP units, whisker movement associated signals were significantly reduced (Fig. 6,7F,H), which suggests that visual cortical silencing has overall a more negative effect on whisker movement related signals in the LP than in the dLGN. Interestingly, whisker movement associated signals were not significantly changed in 62% of both dLGN and LP neurons (Fig. 6,7F,I) indicating that these cells may be modulated by whisker movement independently of the visual cortex. Taken together, these results support the hypothesis that movement-related signals in the majority of visual thalamic cells are not dependent on visual cortical activity and may therefore receive these signals from another brain structure. The 3-5 Hz rhythm on the other hand requires an intact visual cortex, which lends support to the hypothesis that this rhythm may reflect a genuine thalamocortical rhythm.

**Figure 6.**
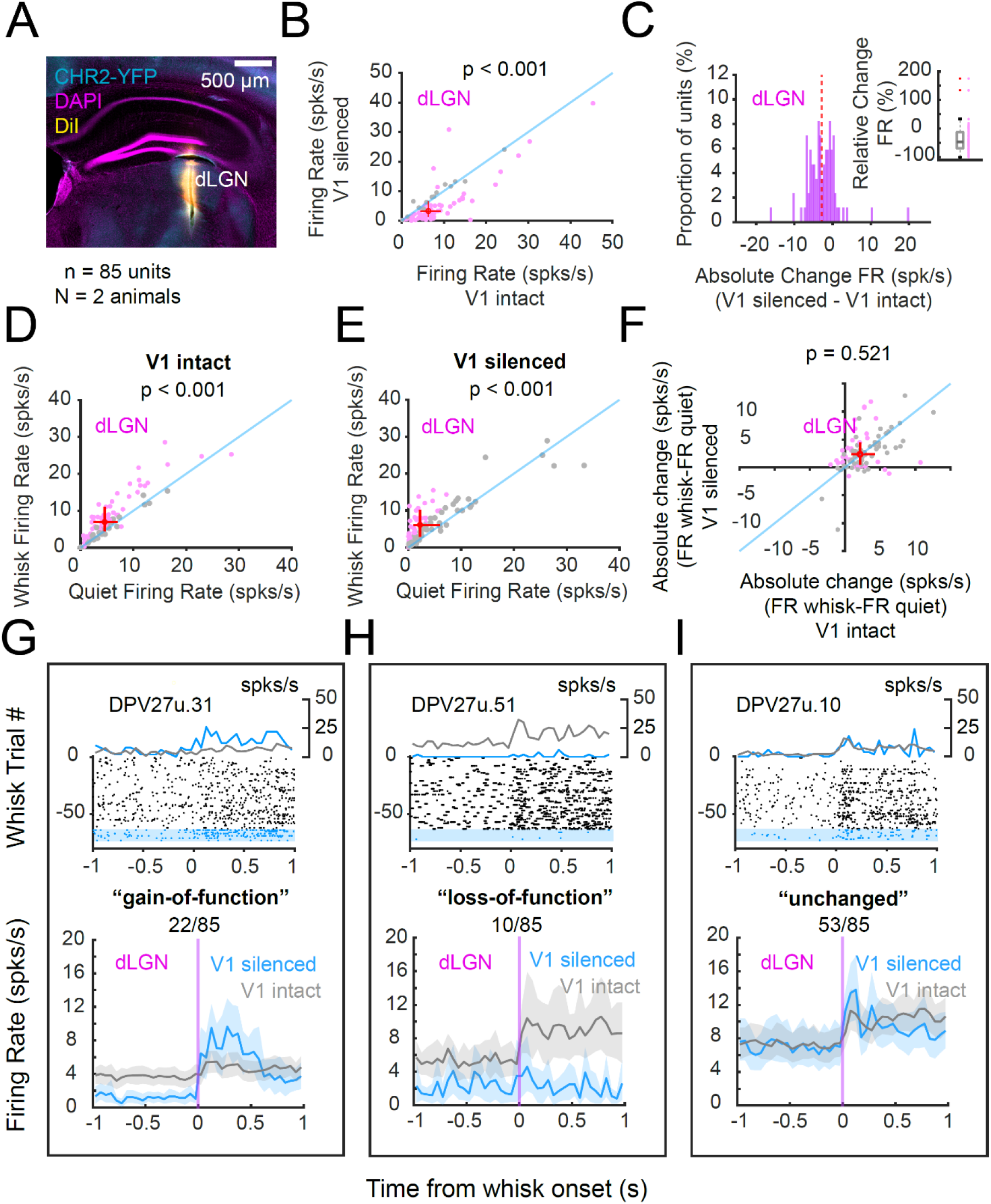
Effects of movement on dLGN thalamic activity with and without silencing of visual cortex. (A) Illustration of the neuropixels recording track demonstrating the recording sites in the medial portion of the dLGN. (B) and (C) Silencing of activity in V1 significantly reduces dLGN activity. Whisker movement is associated with increased dLGN activity whether the visual cortex is intact (D) or inhibited (E). (F) The effects of V1 inactivation on the ability of whisker movement to modulate dLGN activity varied between neurons (pink dots represent units with statistically significant difference between V1 intact and silent). (G-I) Examples and group averages of LP cells that increase (G), decrease (H) or have no significant change (I) in their whisker-related activity with visual cortical inactivation. The paired t-test (for normally distributed data), or the Wilcoxon signed rank sum test (non-parametric) were applied to test statistical significance in D-F. The crosses in scatter plots show the 25th and 75th percentiles with the center representing the overall medians. Data points in F represent the average of bootstrapped means. Statistical significance for each of the data points in F was assessed by testing whether the bootstrapped 95% confidence intervals (not shown) crossed the reference line (shown in blue). Shaded regions in G-I indicate 95% bootstrapped confidence interval. See also Figure S5.

**Figure 7.**
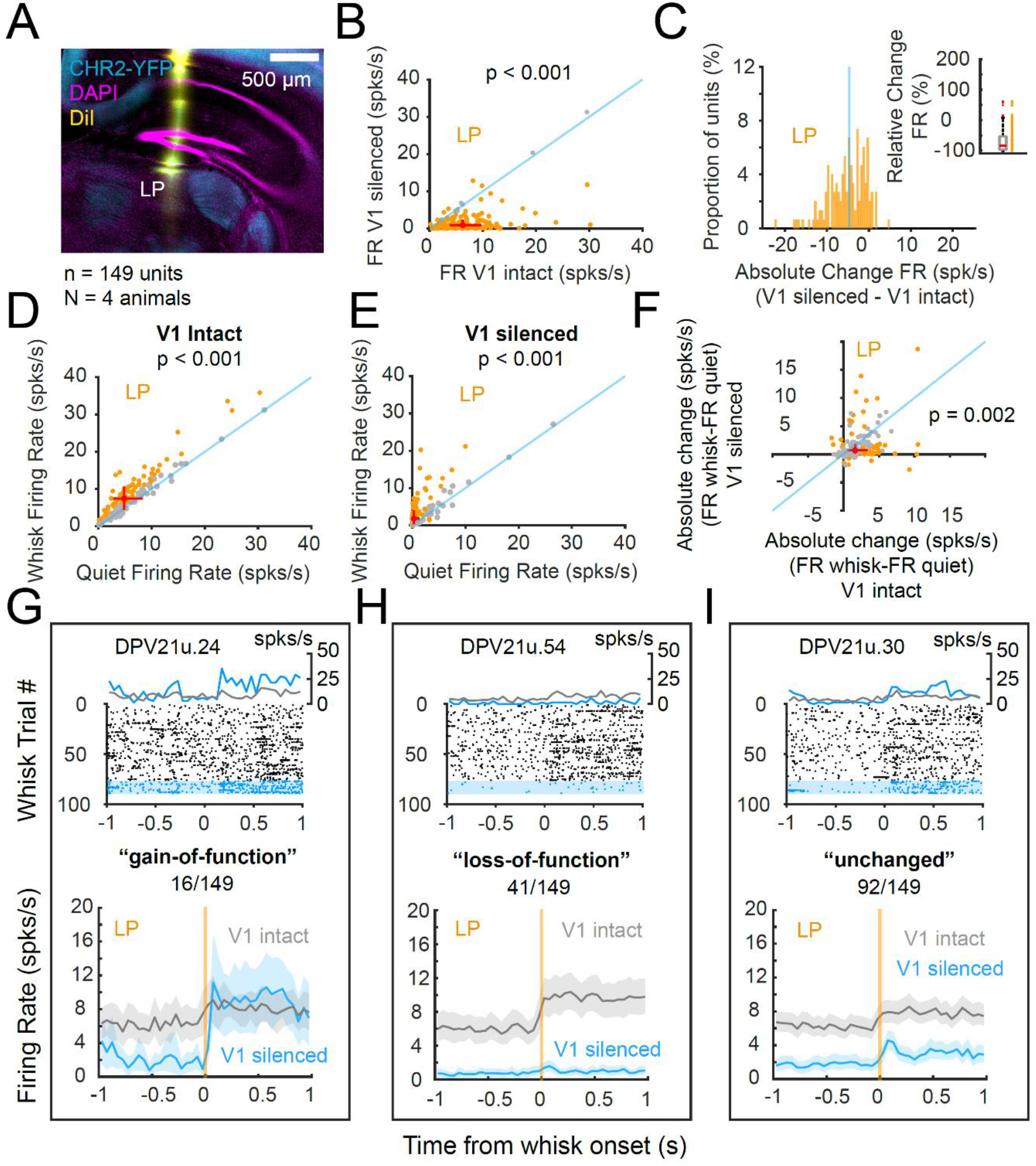
Effects of movement on LP thalamic activity with and without silencing of visual cortex. (A) Illustration of the neuropixels electrode track through the LP nucleus. (B) Silencing of the visual cortex dramatically reduces activity in many LP neurons. (C) Absolute change in firing rate of LP neurons with V1 silencing. (D) and (E) whisker movement significantly increases the firing rate of LP neurons in either normal (D) or visual-cortical silenced (E) conditions. Orange dots represent cells with a significant change in activity with whisker movement vs. quiescence. (F) Distribution of whisker movement induced change in LP firing rate with and without visual cortical silencing (orange dots represent units with statistically significant difference between V1 intact and silent). Examples and group averages of LP cells that increased their whisker movement-associated firing rate changes (G), decreased (H) or showed no significant change (I) with silencing of V1. The paired t-test (for normally distributed data), or the Wilcoxon signed rank sum test (non-parametric) were applied to test statistical significance in D-F. The crosses in scatter plots show the 25th and 75th percentiles with the center representing the overall medians. Data points in F represent the average of bootstrapped means. Statistical significance for each of the data points in F was assessed by testing whether the bootstrapped 95% confidence intervals (not shown) crossed the reference line (shown in blue). Shaded regions in G-I indicate 95% bootstrapped confidence interval. See also Figure S5.

### Pharmacological inactivation of visual thalamus blocks 3-5 Hz rhythm and sustained movement-related depolarizations in V1

To further test the hypothesis that the 3-5 Hz rhythm requires thalamocortical interactions and to test the role of thalamic activity in regulating whisker movement associated signals in visual cortex, we next silenced the visual thalamus and studied state-dependent activity in L2,3 V1 neurons with whole cell recordings. To this end, we inactivated visual thalamus through the injection of fluorescently-coupled muscimol (a potent GABA_A_ receptor agonist), with the center of the injection site in either dLGN or LP, but typically near the border of these two visual thalamic nuclei (Fig. S6). Following silencing of the visual thalamus, we performed intracellular recordings in V1 (Fig. 8). Our results revealed that inactivation of visual thalamic nuclei strongly reduced movement related signals in V1. Specifically, the median amplitude of the rapid PSP responses at whisk onset was significantly reduced to 5.8 mV [7.3-4.7 IQR, unpaired t-test, p = 0.049] which corresponds to a decrease of ~30% when compared to the control cells shown in Fig. 2 (Fig. 8C). The kinetics of the rapid whisker associated depolarizations remained similar compared to controls with a median half-maximum time of 45.1 ms [58.4-13.1 IQR] (Wilcoxon rank sum test, p = 0.146) and a 20%-80% rise time of 229.2 ms [396.9-109.0 IQR] (Wilcoxon rank sum test; p = 0.234) (Fig. 8 E,F). Most strikingly, we observed that the steady-state component of whisker-related signals was almost entirely abolished (median 0.34 mV [0.58-(−0.02) IQR]) when compared to control neurons (median 3.84 mV [5.92-2. IQR] (Wilcoxon rank sum test, p < 0.001) (Fig. 8D). This marked reduction in sustained depolarization with movement following inactivation of the visual thalamus suggests that state-dependent changes of activity in the thalamus are critical to sustain depolarizations associated with arousal/movement related signals in V1.

**Figure 8.**
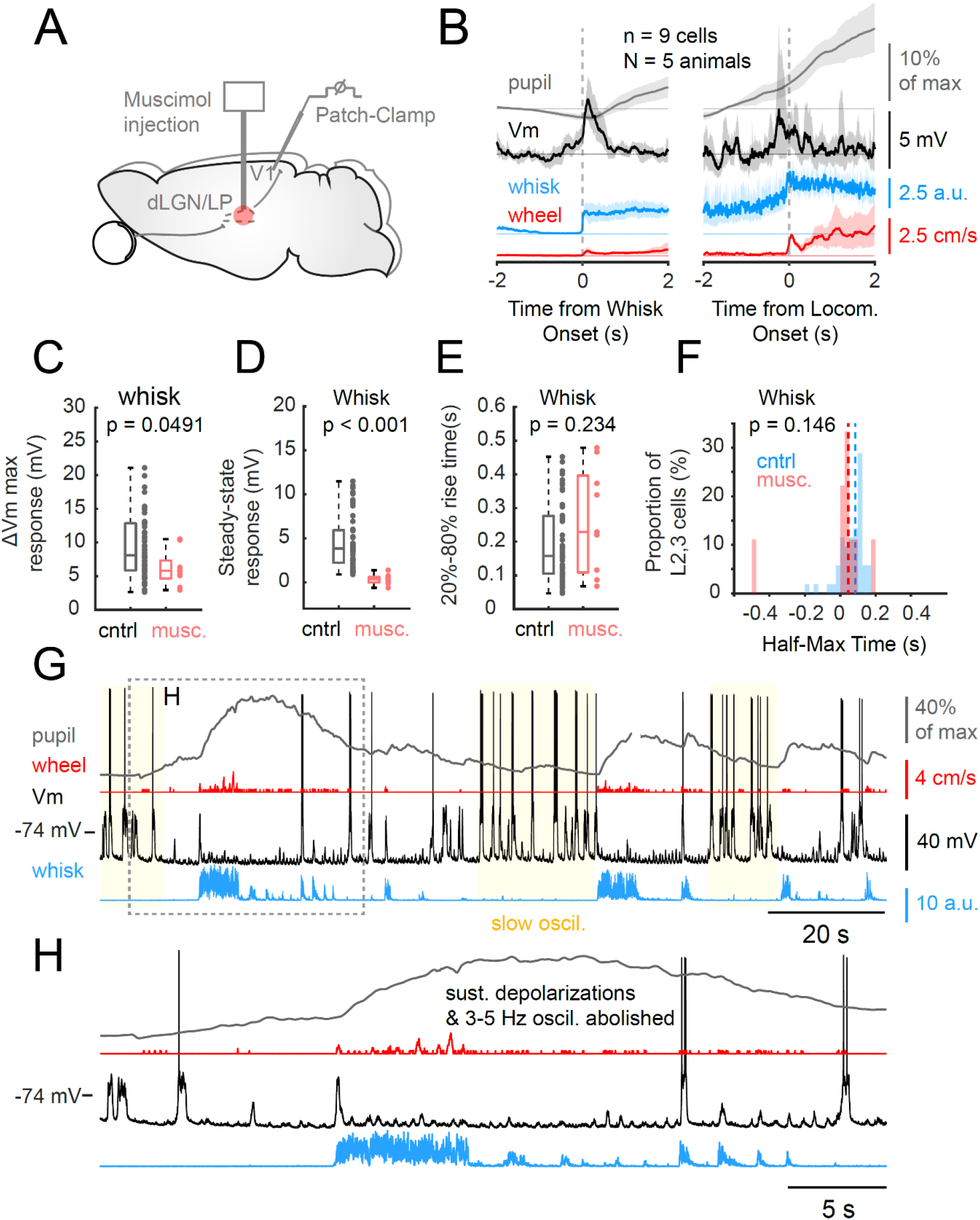
Effects of silencing of visual thalamus on state dependent activity in V1. (A) Illustration of experimental situation. The visual thalamus was silenced with microinjection of the GABAA agonist muscimol. Following injection, whole cell recordings were performed in V1 of awake mice. (B) Alignment of visual cortical neuronal membrane potential with the onset of whisker pad movement or of locomotion revealed transient depolarizations that were not sustained. The depolarization was more highly synchronized with the onset of whisker pad movement than locomotion. The change in membrane potential in cells recorded during silencing of activity in the visual thalamus (n = 9 cells in N = 5 mice) contained a transient component that was significantly smaller than in control cells (controls = same data as shown in Fig. 2) (C), and which exhibited a greatly reduced sustained component during whisker movement (D). (E) The 20%-80% rise time of the transient depolarization was not affected by inhibition of the visual thalamus. (F) Distribution of half-maximum time for control and thalamic-inhibited cells. (G) Example of whole cell recording of a representative L2,3 V1 neuron in the waking mouse following inhibition of the visual thalamus. Note the lack of sustained depolarizations, particularly during bouts of whisker pad movement. (H) Expansion of a portion of the recording in G illustrating the transient nature of synaptic barrages during whisker and wheel movement. See also Figure S6.

In addition to the lack of sustained depolarization of V1 L2,3 neurons during movement following inactivation of the thalamus, we also observed a complete lack of 3-5 Hz oscillations (~84 min of high quality recording time in 9 cells). In control recordings, we would have expected at least 50-75 bouts of 3-5 Hz oscillations during this time period based on the median intervals of one oscillation per 66.67 s and 100 s, that we determined above for LP and dLGN units. This result indicates that the 3-5 Hz oscillation represents a thalamocortical rhythm that depends upon intact interactions between the thalamus and cortex.

## Discussion

In the present study, we investigated the role of visual thalamic activity in regulating waking state-dependent activity in V1, with a particular focus on movement/arousal related signals and a 3-5 Hz alpha-like oscillation. Overall, we found that thalamocortical activity plays an essential role in the generation of the 3-5 Hz oscillations and that thalamic activity is necessary to sustain movement/arousal related signals in V1. Finally, we demonstrate that these signals contain a rapid component that is likely mediated via an ionotropic receptor activating pathway that may act, at least in part, independently of movement, per se.

### 3-5 Hz alpha-like oscillations reflect a thalamocortical rhythm that involves LTS mediated AP firing in visual thalamic neurons

By performing whole-cell recording in L2,3 V1 pyramidal neurons of awake mice, we observed the frequent occurrence of a 3-5 Hz oscillation in the membrane potential (Einstein et al., 2017) (Fig. 1,5). This particular rhythm was previously proposed to constitute an evolutionary precursor of the primate alpha oscillation, and to be associated with pupil constriction, immobility, the offset of visual stimulation and to exhibit the greatest power in LFP in cortical layer 4 (Einstein et al., 2017; Senzai et al., 2019). Our results here demonstrate that this oscillation represents a cyclical thalamocortical interaction, typically associated with decreased movement/arousal, and associated with LTS mediated bursting firing in thalamocortical cells (Fig. 5F-H, Fig. S4). We propose the 3-5 Hz oscillation primarily represents a cyclical thalamocortical interaction based upon: 1) the prevalence of barrages of postsynaptic potentials (PSPs) in both thalamic and cortical neurons with each cycle of the oscillation; 2) the synchronization of these cortical PSPs barrages with AP activity in the thalamus; 3) the initiation of LTSs in thalamic cells by these synaptic barrages (Fig. 5G,H and S4); and 4) the abolition of this oscillation when activity in either visual cortex or thalamus is blocked (Figs. 7,8).

We propose that the 3-5 Hz oscillation is generated through the following events: the withdrawal of a sustained depolarizing synaptic activity - such as after movement, or visual stimulation (or, in primates, upon eye closure), or with low levels of vigilance - leads to a pronounced hyperpolarization of visual thalamic and cortical neurons. The withdrawal of excitatory drive may occur through the failure of corticocortical recurrent drive, such as occurs at the end of an Up state (Compte et al., 2003), through the failure of thalamo-cortico-thalamic interactions to maintain sustained activity, or through the sudden withdrawal of an ionotropic-receptor mediated activating system, such as cholinergic activation mediated through nicotinic receptors (Zagha and McCormick, 2014). The pronounced hyperpolarization removes inactivation of the LTS mechanism in visual thalamic neurons, resulting in a rebound burst of action potentials in a subset of these neurons. This synchronized burst of thalamocortical neuronal activity then drives the visual cortex, resulting in a synchronized discharge of visual cortical neurons, including thalamic projecting layer 5/6 neurons, which then initiate a new cycle of activity within the visual thalamus. The oscillation continues, with each cycle weakening in relation to a decreased intercycle hyperpolarization in thalamocortical neurons, perhaps owing to h-current activation (Lüthi and McCormick, 1998), synaptic depression (Chung et al., 2002; Nestvogel et al., 2020), or desynchronization of thalamocortical activity (Wright et al.).

The thalamic reticular nucleus (TRN) plays a prominent role in spindle waves during sleep and high-threshold spindles in waking rats (Buzsaki et al., 1988; Fernandez and Lüthi, 2020; McCormick and Bal, 1997; Steriade et al., 1993). The role of the thalamic reticular nucleus in the 3-5 Hz oscillation studied here is as of yet unknown. The possibility that the hyperpolarizing periods in thalamocortical neurons are initiated by activation of TRN neurons can be examined in future studies by simultaneous recording of TRN and thalamocortical neurons and determining the reversal potential of the hyperpolarizing and depolarizing cycles of the oscillation in thalamocortical neurons.

How does this model compare to the results of previous studies? With the increase in brain size, it was previously proposed that the frequency of the alpha rhythm may have increased from 3-5 Hz in the mouse, to 6-12 Hz in cats and to 8-12 Hz in humans (Senzai et al., 2019). It is not clear whether the neural mechanisms that generate the alpha oscillation were conserved in evolution. In cats, it has been proposed that high-threshold mediated AP bursts are associated with alpha rhythms (Crunelli et al., 2018; Hughes and Crunelli, 2005; Hughes et al., 2004; Lorincz et al., 2009), which is different to our results of LTS mediated AP bursting during the 3-5 Hz alpha-like rhythm in the mouse. Interestingly, mice and primates are closer relatives in evolutionary terms than cats and primates, and it remains to be addressed, whether LTS, or high threshold mediated AP bursts are associated with the primate alpha oscillation and whether the degree by which animals are related to each other in evolutionary terms, or the size of the brain is the decisive factor that determines the mechanism of alpha rhythm generation. Although the cell intrinsic mechanisms that are associated with the alpha rhythm may differ in the animal kingdom, it is intriguing to speculate that the overall importance of visual thalamic activity in regulating the alpha rhythm in visual cortical regions may have been conserved across mice, cats and primates (Hughes and Crunelli, 2005; Saalmann et al., 2012). Interestingly, recent studies in mice suggest that the presence of the 3-5 Hz rhythm is disruptive to performing a learned Go/No-Go response to a visual stimulus (Speed et al., 2019), suggesting that it may occur mainly during periods of inattentiveness or may, itself, actively disrupt thalamocortical sensory processing.

### Sustained depolarizations in V1 neurons during movement/arousal correlate with, and require, visual thalamic activity

The second state-dependent activity pattern that we focused on in our study was the rapid change from slowly fluctuating synaptic activity to a steady depolarisation in V1 neurons that has been observed in conjunction with locomotion and increases in arousal (Bennett et al., 2013; Polack et al., 2013; Reimer et al., 2014) (Fig. 1). This distinct brain state has also been reported in the primary auditory cortex (A1) (McGinley et al., 2015a; Schneider et al., 2014) and somatosensory cortex (S1) (Poulet and Petersen, 2008; Zagha et al., 2013). Various neural mechanisms have been proposed to regulate this state-dependent activity change including neuromodulation via noradrenergic and cholinergic pathways (Eggermann et al., 2014; Fu et al., 2014; Lee et al., 2014; Polack et al., 2013), changes in the activity of intracortical inhibitory networks (Fu et al., 2014; Polack et al., 2013; Reimer et al., 2014), and feedback from secondary and primary motor cortex (Leinweber et al., 2017; Zagha et al., 2013). In addition, studies have shown that inactivation of the somatosensory thalamus abolishes the steady depolarisation in S1 neurons (Poulet et al., 2012), suggesting that thalamic activity may provide the synaptic input necessary for the sustained depolarization in the cortex.

Here we performed a detailed kinetic analysis in L2,3 V1 neurons of the sustained depolarization associated with movement/arousal. Our results revealed a tight coupling of whisker movements with the occurrence of these depolarizations on the scale of tens of milliseconds or less (Fig. 2), lending support to the hypothesis that these responses are driven in V1 by fast ionotropic signalling. Importantly, similar depolarizations also occurred during pupil dilations in the absence of overt movements of the whisker pad and the running wheel (Fig. S1), suggesting that a significant component of these signals may be associated with changes in arousal rather than movement per se (McGinley et al., 2015a; Petty et al., 2021), with the neural mechanisms of movement adding additional activating and information-containing (e.g. eye position, head direction) signals (Bouvier et al., 2020; Miura and Scanziani, 2021).

Simultaneous extracellular recordings in visual thalamic neurons with whole-cell recordings in V1 additionally revealed that the rapid depolarizations in V1 strongly correlated with visual thalamic activity (Fig. 3). Interestingly, we also found that the membrane potential of visual thalamic neurons exhibited depolarizations with similar kinetics as those of V1 neurons during whisk onsets indicating these two brain structures exhibit remarkable similarities in movement/arousal related state changes on the level of AP firing and postsynaptic activity (Fig. 4). The membrane potential of thalamic neurons is highly regulated by a variety of modulatory neurotransmitters (McCormick, 1992). Currently, the only known modulatory system that can quickly depolarize thalamic neurons is cholinergic activation of nicotinic receptors and corticothalamic activation of glutamatergic ionotropic receptors (McCormick, 1992), although the possible involvement of an additional ionotropic excitatory releasing neurotransmitter system in the observed movement-related thalamic activations cannot be excluded. Relevant to this, recent work suggests that neither reafferent signals from the periphery, the superior colliculus, or cerebral cortex are responsible for state-dependent changes in sensory thalamic activity (Petty et al.), although state changes in retinal activity (Liang et al., 2020; Schröder et al., 2020) may contribute to state-dependent activity in the visual thalamus.

In contrast to the lack of consistent effect of visual cortical silencing on arousal/movement related changes in the visual thalamus (Fig. 6,7), silencing of the visual thalamus dramatically affected visual cortical activity by largely abolishing the movement-related steady depolarization. Following inhibition of thalamic activity, the only movement-related component remaining in the visual cortex was a transient depolarization at movement onset, which was also reduced in amplitude, but not kinetics (Fig. 8). These results indicate that thalamic activity plays a critical role in sustaining movement/arousal related signals in V1, and that there may be additional ionotropic pathways mediating the rapid, transient component.

These results are very similar to the findings of Poulet et al in S1 (Poulet et al., 2012), except for the presence of the residual rapid depolarization at the onset of whisking in our study. We speculate that the transient onset may originate from cortical input by structures such as the primary and secondary motor cortices (Leinweber et al., 2017; Schneider et al., 2014; Zagha et al., 2013), or alternatively from fast nicotinic signalling via cholinergic input from the basal forebrain (Eggermann et al., 2014; Reimer et al., 2016), which in turn may involve the action of brain stem nuclei such as the mesencephalic locomotor region (Lee et al., 2014). Given that our pharmacological injections only partially inactivated the visual thalamus, as indicated by the histological assessment of fluorescent signals of the injection sites (Fig. S6), we cannot rule out the possibility that the residual signal at the onset of the depolarization in V1 during our experiments could also originate from the activity of visual thalamic neurons that were not silenced. Future studies are required to test these possibilities in greater detail, which in turn may help to better understand the specific contribution of the other previously proposed neural mechanisms to regulating and shaping the movement/arousal related state-changes in V1 and other primary sensory cortices described above (Eggermann et al., 2014; Fu et al., 2014; Leinweber et al., 2017; Polack et al., 2013; Zagha et al., 2013). Based on previous results of the necessity of activity of the somatosensory thalamus for whisker movement related signals in S1 and our similar results in the visual system, we speculate that thalamocortical interactions play an essential mechanistic role in regulating movement/arousal related cortical state changes that is conserved across primary sensory regions.

In summary, our study supports the hypothesis of a crucial role of visual thalamocortical interactions in regulating and shaping reccurent waking state-dependent activity. These state-dependent changes, in turn, have been reported to significantly contribute to the astonishing trial-to-trial variability of sensory and behavioral responses (McCormick et al., 2020; McGinley et al., 2015a, 2015b; Neske et al., 2019; Niell and Stryker, 2010; Speed et al., 2019).

## Acknowledgements

We thank Cristopher Niell and Daniel Hulsey for helpful comments on the manuscript and Rennie Kendrick and Kevin Zumwalt for technical assistance. We also thank Laura Boddington, Lindsay Collins, Suh Yun Jo, Paul Steffan and Evan Vickers for helpful discussions. This work was supported by NIH grants R35NS097287 and R01NS118461.

## Author Contributions

D.B.N. and D.A.M. conceived and designed the study. D.B.N. performed experiments and data analysis. D.B.N. and D.A.M. wrote the paper.

## Methods

### In-vivo Patch Clamp Recordings

Blind, in vivo whole-cell recordings were performed in the primary visual cortex of adult mice (> 2 months). For this, a craniotomy (~300 μm diameter) was introduced under isoflurane anesthesia at 3 mm posterior of bregma and 2.5 mm lateral to the midline (Lien and Scanziani, 2013). Mice were allowed to recover from surgery for at least 3 hours, before recording. Patch pipettes (4-7 MΩ) were inserted into the brain with an initial positive pressure of 300-400 mbar (for V1 recordings), or at 800 mbar (for thalamic recordings). The pressure was reduced to 30 mbar, upon reaching a depth of ~100 μm for V1 recordings (~2300 μm for thalamic recordings). To search for neurons, the pipette was lowered at 2 μm steps and increases in the pipette resistance were monitored via the application of a test pulse in voltage clamp mode (Margrie et al., 2002). V1 neurons were measured at a depth of 100-350 μm and thalamic cells at a depth of 2300-2700 μm from the pial surface. Successful encounters with cells were followed by the formation of a giga seal at a holding potential of −70 mV and the cell membrane was ruptured via short negative pressure pulses to perform recordings in the whole-cell/current clamp mode (series resistance < 60 mΩ) (Hamill et al., 1981). The intracellular recording solution contained (in mM) 130 K gluconate, 4 KCl, 2 NaCl, 10 HEPES, 0.2 EGTA, 4 ATP-Mg, 0.3 GTP-Na, and 14 phosphocreatine-2K and 2% biocytin adjusted to a pH of 7.4 and at an osmolarity of ~290 mM. The surface of the brain was kept moist during recordings with Ringer’s solution (in mM: 145 NaCl, 5 KCl, 1.8 CaCl_2_, 1 MgCl_2_, 5 HEPES adjusted to pH 7.4, ∼290 mM osmolarity). All whole-cell recordings were carried out using a multiclamp 700B amplifier (Molecular Devices) and signals were sampled at 50 kHz using a Power 1401 digitizer (Cambridge Electronic Design Limited). Putative excitatory neurons were distinguished from inhibitory neurons by their relatively broad AP half-width and a regular firing pattern upon positive current injections (McCormick et al., 1985). The initial recording period, in which the membrane potential was significantly hyperpolarized was excluded from the analysis (McGinley et al., 2015a) and only neurons with more than 400 s of high quality recording time (stable membrane potential, overshooting APs during recording) were included in the analysis. The resting membrane potential was corrected offline for a liquid junction potential of −14 mV (Cruikshank et al., 2007). APs were detected when dV/dt reached a threshold of 25 V/s. For subsequent analyses of postsynaptic potentials during state changes, all APs were removed from the recordings by using a custom-written Matlab script (time window of −1x AP halfwidth + 3.5x AP halfwidth) (Nestvogel et al., 2020). The maximum of PSP responses associated with whisker movement and locomotion was identified within a window of −0.5 to +0.5 s from movement onset. The 20% to 80% rise time was identified within a time window of −0.5 to +0.5 s from the peak PSP response. The average steady-state amplitude of depolarizations associated with whisker movement and locomotion were calculated within a time window of 0.5-2 s after motor movement onset.

### Extracellular Recordings

Extracellular recordings were made with Neuropixels probes (neuropixels 1.0, a.k.a. phase 3b)(Jun et al., 2017). Electrode shanks were coated in Dil or DiO (Vybrant solution, Thermo Fisher Scientific) before recordings to allow the post hoc identification of the recording track. A small craniotomy (~300 μm) was introduced under isoflurane anesthesia at 2.0-2.1 mm posterior of bregma and 2 mm lateral to the midline (for LP recordings), or at 2.5 mm posterior of bregma and 2.3 mm lateral to midline (for dLGN recordings). Mice were allowed to recover for at least 3 hours before electrophysiological recordings began. The surface of the exposed brain was kept moist during experiments with Ringer’s solution as described above. The electrode shanks were lowered with a Sutter Instruments micromanipulator at low speed (~2-3 μm per seconds) until a depth of 3600-4000 μm was reached. Recordings were performed after a minimum of 15 min of reaching the final depth at a gain of 250 (LFP), and 500 (APs) with the open-ephys software. All extracellular recording data was acquired using a PXIe acquisition module by National Instruments. A sync signal was sent from the acquisition module to the Power 1401 digitizer in order to synchronize neuropixels data with patch clamp recordings and camera frames. The neuropixels data was sampled at a setting of 30 kHz, but we detected slight deviations from this number offline (~ e.g. actual rate for one recording 29999.795 Hz). All recordings were analyzed at the actual sampling rate. Thalamic AP burst firing was detected with interspike intervals of less than 4 ms (>250 Hz) preceded by more than 100 ms of AP quiescence.

### Videography and Movement Analysis

The anterior half of the animal’s body was monitored via videography at a frame rate of 125 Hz using a FLIR Grasshopper 3 camera. Images were acquired with the FlyCapture2 software (FLIR). A digital trigger preceded the acquisition of each frame and was accomplished using a Power 1401 digitizer. The body and the eyes were illuminated by infrared light, which was present in addition to a constant ambient light in the room. The area of the pupil was determined offline using FaceMap (Stringer et al., 2019), or DeepLabCut (Nath et al., 2019). The motion energy of whisker pad movements was calculated with a custom matlab script by subtracting the sum of the absolute pixel intensity of each frame from that of the preceding frame. A whisker movement-bout was defined by the normalized motion energy (to maximum) reaching a threshold of 1 (a.u.) and staying above the threshold for at least 2 seconds. Neighboring whisker movement-bouts that occurred within 1 s or less were grouped together. Similarly, locomotion was defined by the wheel velocity reaching a threshold of 1 cm/s and lasting for at least 1 s. Neighboring locomotion-bouts that occurred within 1 s or less were grouped together.

### Immunohistochemistry and Microscopy

Following electrophysiological recordings, mice were deeply anesthetized and subjected to whole-animal perfusion using phosphate buffer (PB) and PB supplemented with 4% paraformaldehyde (PFA). Brains were removed afterward and placed in 4% PFA for at least one night. Brain slices (200 μm thickness) were prepared by using a cryostat after brains were washed 3 times with PB and after being placed consecutively in 20% and 30% sucrose solution. Before immunostainings, brain slices were incubated at room temperature in 3% Triton-X PB solution. The brain slices were then transferred into a 3% Triton-X PB solution supplemented with Alexa Fluor 555 streptavidin (1:1000, Thermo Fisher Scientific) and DAPI (1:1000, Thermo Fisher Scientific) and incubated at room temperature for an additional 4 hours. In the last step, brain slices were washed 2-3 times with PB (20-30 min) and mounted onto glass microscope slides for subsequent imaging. Microscopy was performed using a CSU-W1 SORA spinning disk microscope (Nikon) equipped with a 4x and a 60x (water) objective.

### Pharmacological and Optogenetic Silencing of dLGN and V1

A volume of 300 nL fluorophore-coupled muscimol (Bodipy, Thermo Fisher Scientific) was injected into the visual thalamus (2400-2600 μm depth; 2.0-2.3 mm lateral to midline and 2.0-2.5 mm posterior to bregma) at a rate of 60 nL/min at a concentration of 2 mM using a Nanoliter 2020 system (WPI). Optogenetic silencing of V1 was achieved by the use of mice in which channelrhodopsin-2 expression was driven by *cre* in parvalbumin positive interneurons (Ai32 x PVcre). A craniotomy with a diameter of ~300 μm was introduced above V1 (2.5 mm lateral to midline and 3 mm posterior to bregma), while mice were under isoflurane anesthesia. During the electrophysiology recordings, an optical monofiber (0.63 NA) was positioned in close proximity to the craniotomy (1 mm above the dura) and light pulses at 9 mW (at fiber tip) and a duration of 8 s were applied (LEDD1B- T-Cube LED driver, Thorlabs; Connectorized LED 465 nm, Doric Lenses) under the control of CED Spike2 software. Single light pulses were separated by 4 s and 30 s to study the impact of silencing V1 activity on state-dependent activity in the LP and dLGN. Although our optogenetic stimulation was centered around V1, we do not rule out the possibility that other areas surrounding V1 may have partially been stimulated, for which reason we refer to this manipulation as a silencing of the visual cortex. We excluded the first 2 s after optogenetic stimulus onset in our data analysis, as well as the first 2 s following the stimulus offset in order to avoid contamination of neural activity associated with stimulus transients.

### Analysis of data obtained with neuropixels probes

All extracellular data obtained with neuropixels were preprocessed using common-average referencing (https://github.com/cortex-lab/spikes/tree/master/preprocessing)(Steinmetz et al., 2019). After preprocessing, spikes were sorted with Kilosort2 (https://github.com/MouseLand/Kilosort) (Pachitariu et al. 2016). Putative single units that were identified by kilosort2 were subjected to manual curating with phy2. During manual curation, each unit was inspected for potential refractory contaminations, robustness of waveform during recording, similarity of waveform to other units and sudden loss of spiking events during recording. Those units, which passed the quality assessment were labeled as single units and were included in the analysis. Only units with an average spike rate of greater than 0.2 spikes/s were included in the analysis. The depth of neuropixel probes was later matched with the microscopic confocal images to identify dLGN and LP units. For this, we first identified the border between the hippocampus CA3 region and the visual thalamus in our electrophysiology recordings, which was characterised by a large gap of no unit activity. By using the scalable brain atlas by the Allen Institute (Allen Mouse brain common coordinate framework version 3) as an aid, we next measured the length of the DiI-stained track spanning the corresponding visual thalamic region in the microscopy images and related this information to the depth of electrophysiological recordings to identify the border between visual thalamic regions and neighboring deeper regions (e.g. PoM). Due to tissue shrinkage during PFA fixation, we assumed that this approach would exclude some units that laid in deeper regions of visual thalamic regions. To determine the spectral coherence between the membrane potential of V1 neurons and visual thalamic activity (average activity of all units per recording), we made use of a Matlab script (Kramer, 2013) with an overlap of 90% and a sliding window of 20s.

### Statistical analysis

All data was tested for normality using the Lilliefors test in Matlab 2019b. For normally distributed datasets we employed the paired t-test (ttest in Matlab), or the unpaired t-test (ttest2 in Matlab) where it was appropriate. For non-parametric data sets, we used the Wilcoxon signed rank sum test (signrank in Matlab), or the Wilcoxon rank sum test (ranksum in Matlab). For multiple comparisons, we made use of the Kruskal-Wallis test (kruskalwallis in Matlab) together with Bonferroni correction (multcompare, ‘CType’, ‘bonferroni’ in Matlab). All 95% confidence intervals shown in graphs were obtained by bootstrapping. To obtain 95% bootstrapped confidence intervals, we randomly resampled the raw data with replacement to calculate a bootstrapped group of means (n = 1000). The confidence intervals reflect 95% of this bootstrapped group. Bootstrapping was also performed for Fig 6,7 F to perform statistical testing on datasets. Differences were deemed to be significant, when the 95% bootstrapped confidence intervals did not cross the reference line (m = 1) in the corresponding plots. The interquartile ranges (IQR) in text and figures reflect the 25th and 75th percentiles (Matlab function: prctile(x,[25 75],‘all’).

---

**Figure S1 (related to Figure 2).**
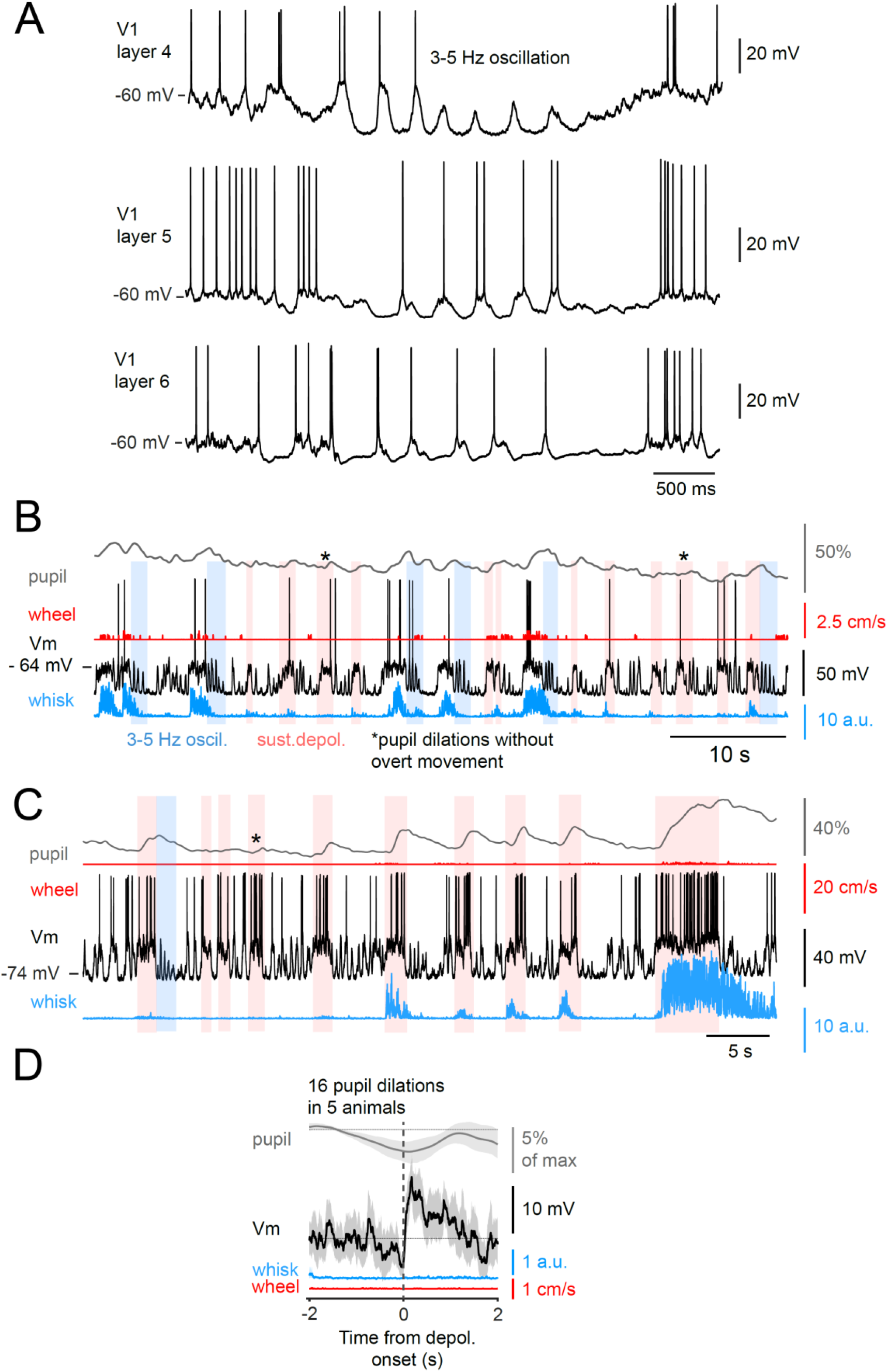
Pupil dilations in absence of overt movements of the whiskers or running wheel are associated with rapid depolarizations in L2,3 V1 neurons and 3-5 Hz oscillations can also be found in neurons of layers 4,5 and 6. (A) Example recordings of layer 4, 5 and 6 neurons illustrating the occurrence of the 3-5 Hz oscillation. (B) and (C) Example recordings of two different L2,3 V1 neurons in which pupil dilations occurred without overt movements of the whiskers or running wheel (which detects not only rotation, but also body postural changes; see Methods). A. is the same cell as shown in Figure S2A and S6F (top) (D) Examination of the membrane potential of 15 randomly selected V1 L2,3 recordings showed that a total of 16 pupil dilations without discernible movements could be found in 5 mice. The figure shows the averaged pupil area, whisker motion energy and locomotion speed sorted according to the rapid depolarization that preceded the pupil dilations. Shaded regions in the graphs indicate the 95% bootstrapped confidence interval.

**Figure S2 (related to Figure 2).**
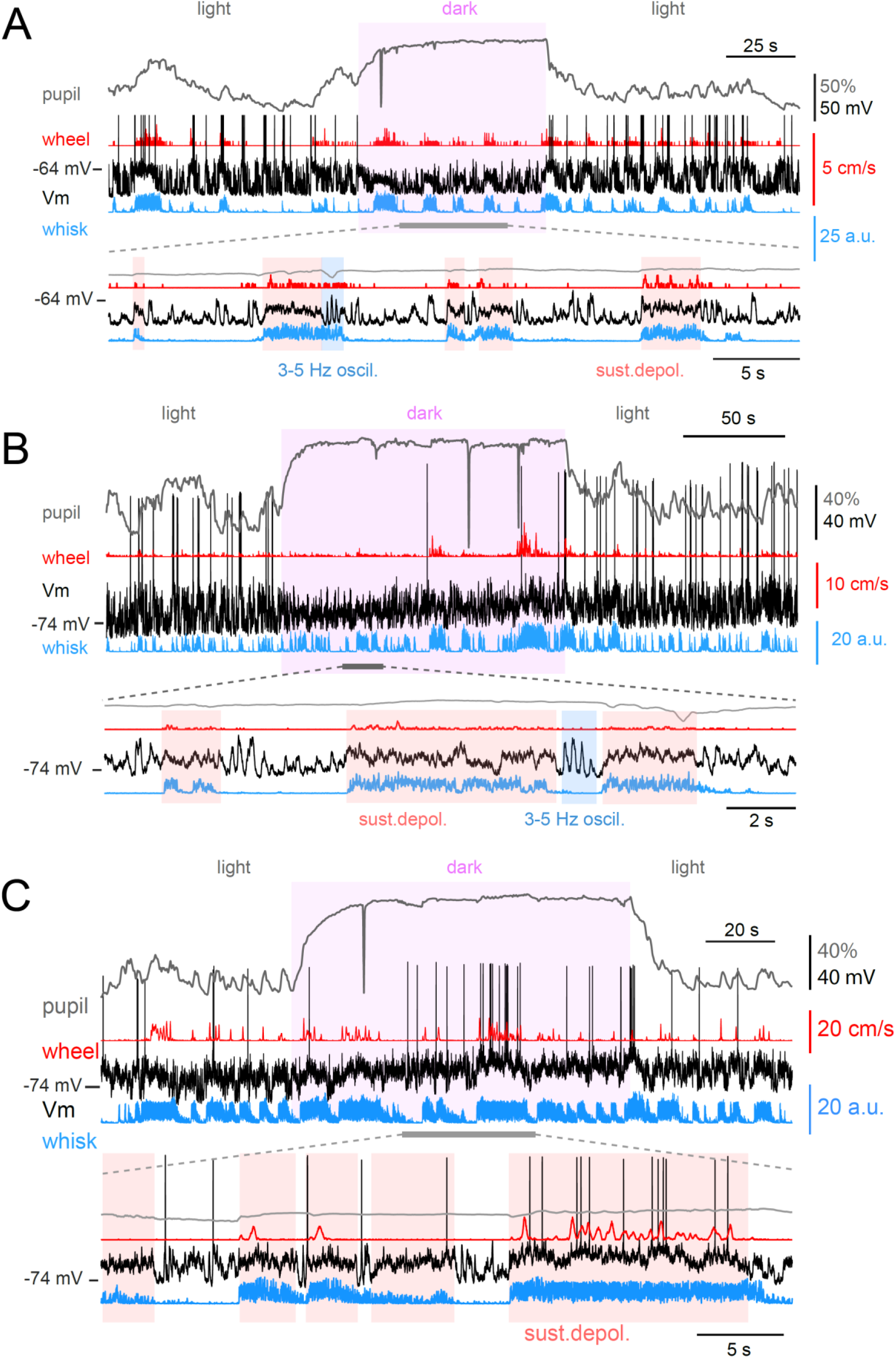
Rapid movement related depolarizations in L2,3 V1 neurons persist in the absence of visible light. (A-C). Example recordings of three different L2,3 V1 neurons with and without visible light. The infrared light was not switched off during these recordings to allow the quantification of whisker pad movements and change in pupil area. Note that in all three cells, depolarizations still occur in the absence of visible light. The neurons shown in A and B exhibited a pronounced reduction in AP firing when the visible light was switched off (purple window). A. is the same cell as shown in Figure S1A and S6F (top)

**Figure S3 (related to Figure 2).**
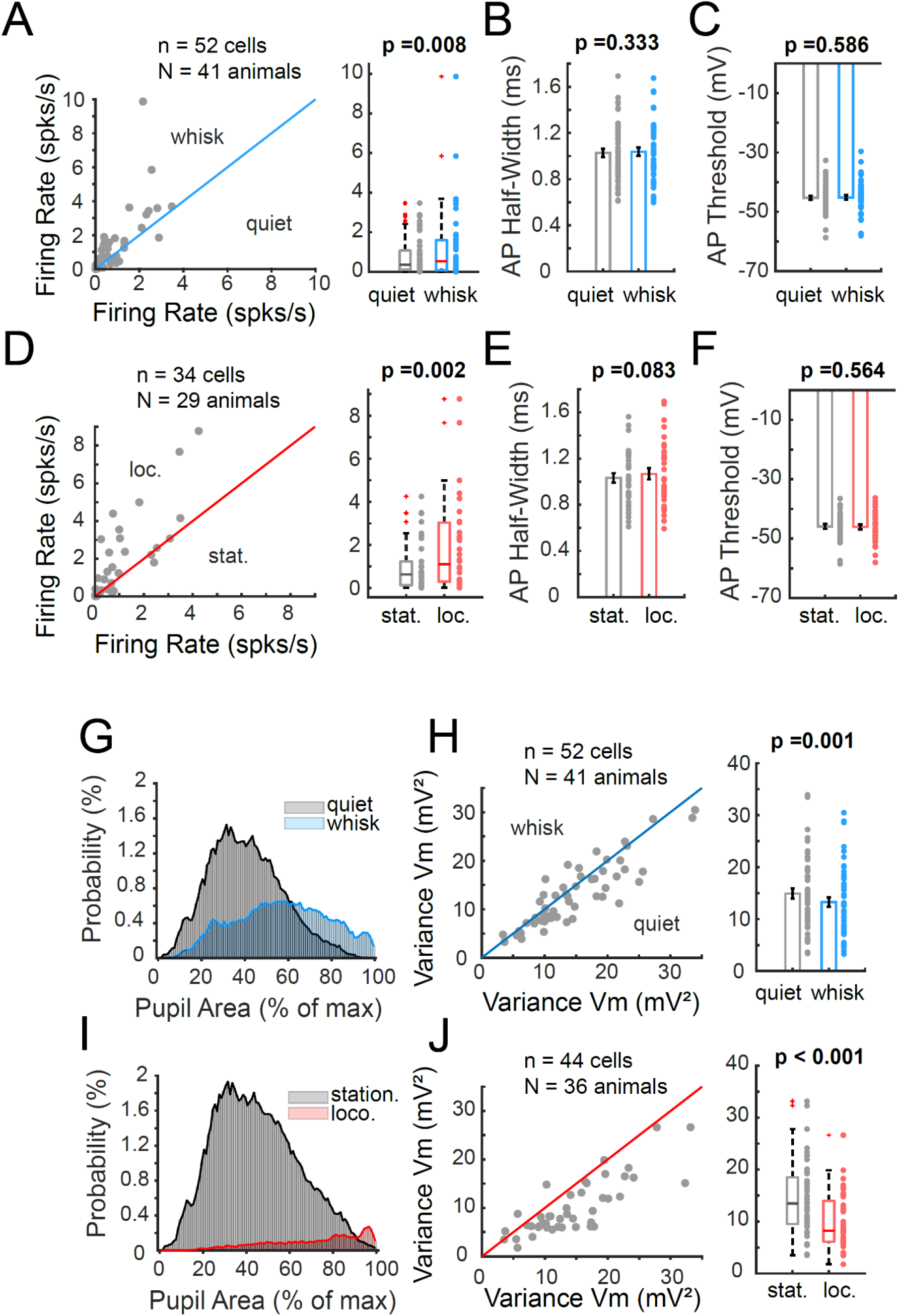
Both whisker movement, and walking with whisker movement, result in an overall increase in firing rate of V1 L2,3 neurons, without a significant change in AP half width or AP threshold. (A) Whisker movement increased the AP activity of a subset of L2,3 V1 neurons recorded intracellularly. Distribution of action potential firing rates in quiescence and whisker movement is shown on the right. AP width at half maximum (B) and threshold (C) are not significantly affected by whisker movement. (D) Locomotion is associated with an increase in firing rate of intracellularly recorded visual cortical neurons. This increase in activity is not associated with a significant change in action potential width (E) or threshold (F). Both whisker movement (G) and locomotion (I) are associated with a shift of pupil diameter towards larger amplitudes. This is particularly true after 1-2 seconds from movement onset (not shown). Data shown here are not adjusted for this pupil diameter lag. Both whisker movement (H) and locomotion (J) are associated with a significant decrease in membrane potential variance in visual cortical neurons. Non parametric datasets are depicted as boxplots which show the median, 25th and 75th percentiles and the whiskers indicate the most extreme data points that are not considered outliers. Normally distributed datasets are depicted as bars and the error bars show the standard error of the mean. The unpaired t-test (for normally distributed data), or the Wilcoxon rank sum test (non-parametric) were applied to test statistical significance.

**Figure S4 (related to Figure 4).**
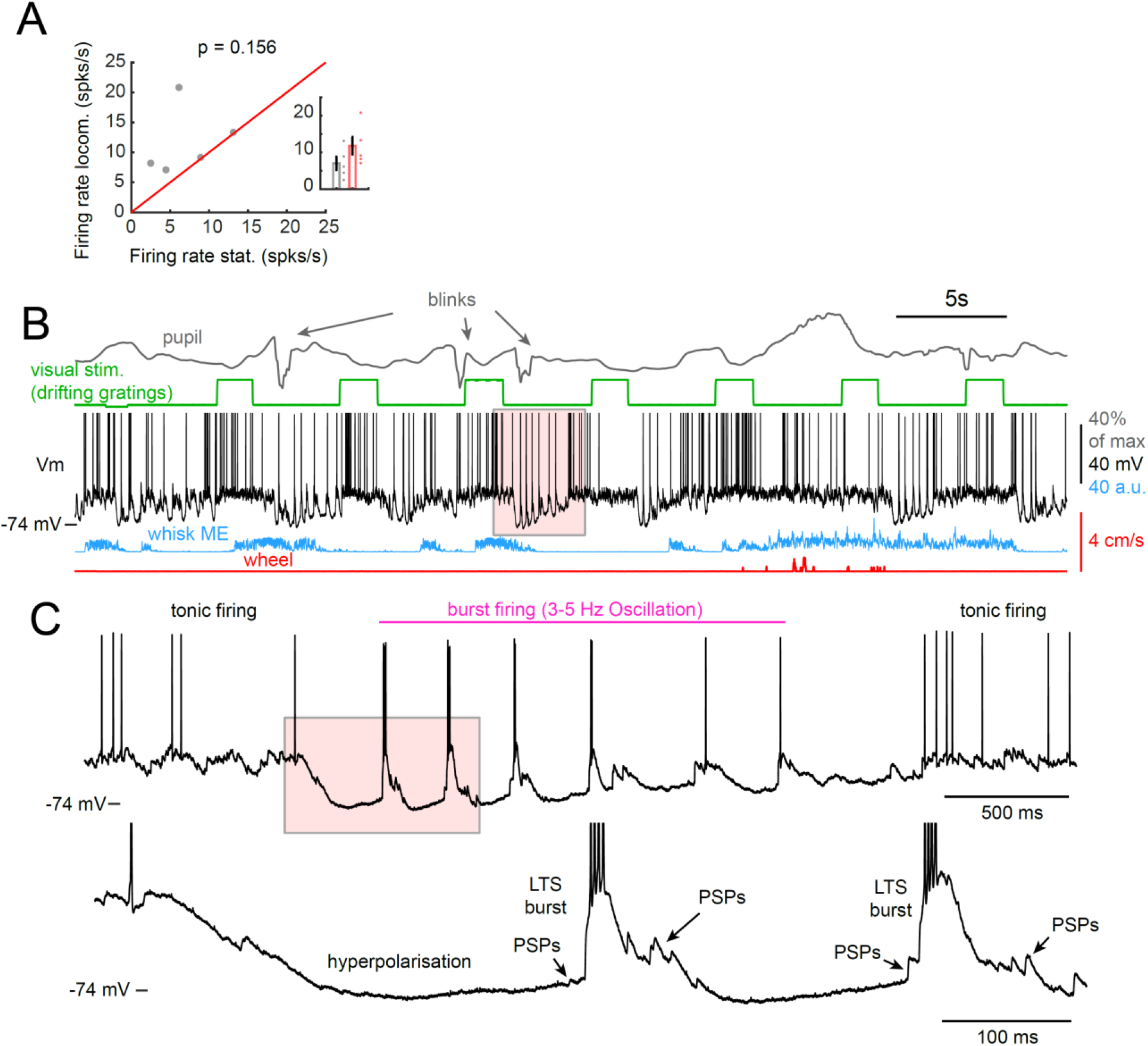
Whole-cell recordings in visual thalamic neurons reveal enhanced AP firing during whisker movements in some visual thalamic neurons and 3-5 Hz oscillations to also occur often after visual stimulation. (A) The AP firing rate is enhanced in 3 out of 5 intracellularly recorded visual thalamic neurons during locomotion (no locomotion bouts occurred in 3 out of the 8 recordings). (B) Example recording of a visual thalamic neuron during visual stimulation. Previous studies showed that the likelihood of the 3-5 Hz rhythm is enhanced with visual stimulation (Einstein et al., 2017). Note that the example visual thalamic neuron also exhibited pronounced LTS mediated burst firing at a 3-5 Hz rhythm, which indicates that the cell intrinsic mechanism for the generation of the 3-5 Hz rhythm is identical between those occurring after visual stimulation offset and those occurring preferentially after enhanced movement activity. (B) Expansion of a 3-5 Hz oscillation occurring after the offset of visual stimulation (red box in A).

**Figure S5 (related to Figures 6,7).**
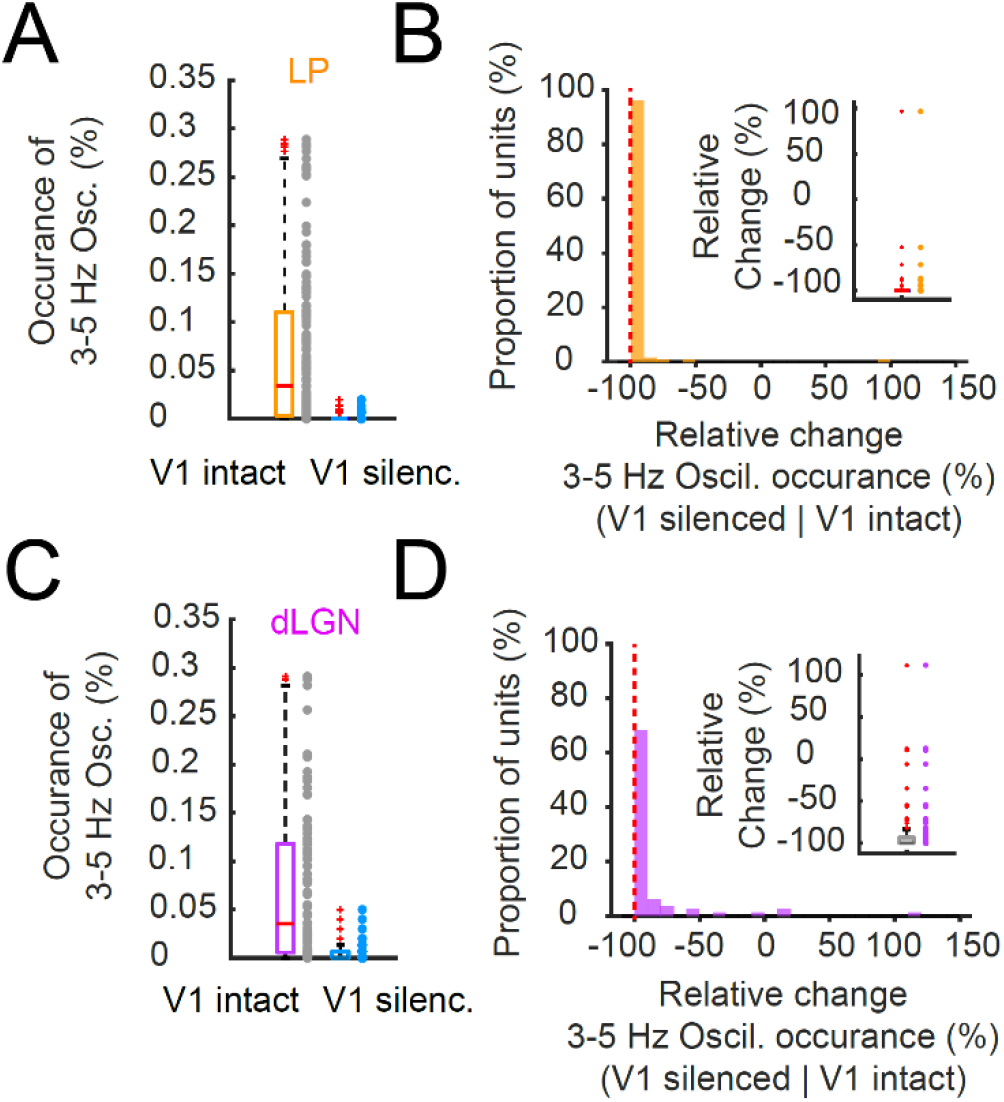
Silencing V1 abolishes the 3-5 Hz oscillation in visual thalamus. Rate of occurrence of 3-5 Hz oscillations in the LP (A) and dLGN (C) with and without visual cortical silencing (including cells with a 0% occurrence). Relative change in rate of occurrence of 3-5 Hz oscillation in LP (B) and dLGN (D) with visual cortical silencing. Note that visual cortical silencing nearly completely abolishes the occurrence of 3-5 Hz oscillations.

**Figure S6 (related to Figure 8).**
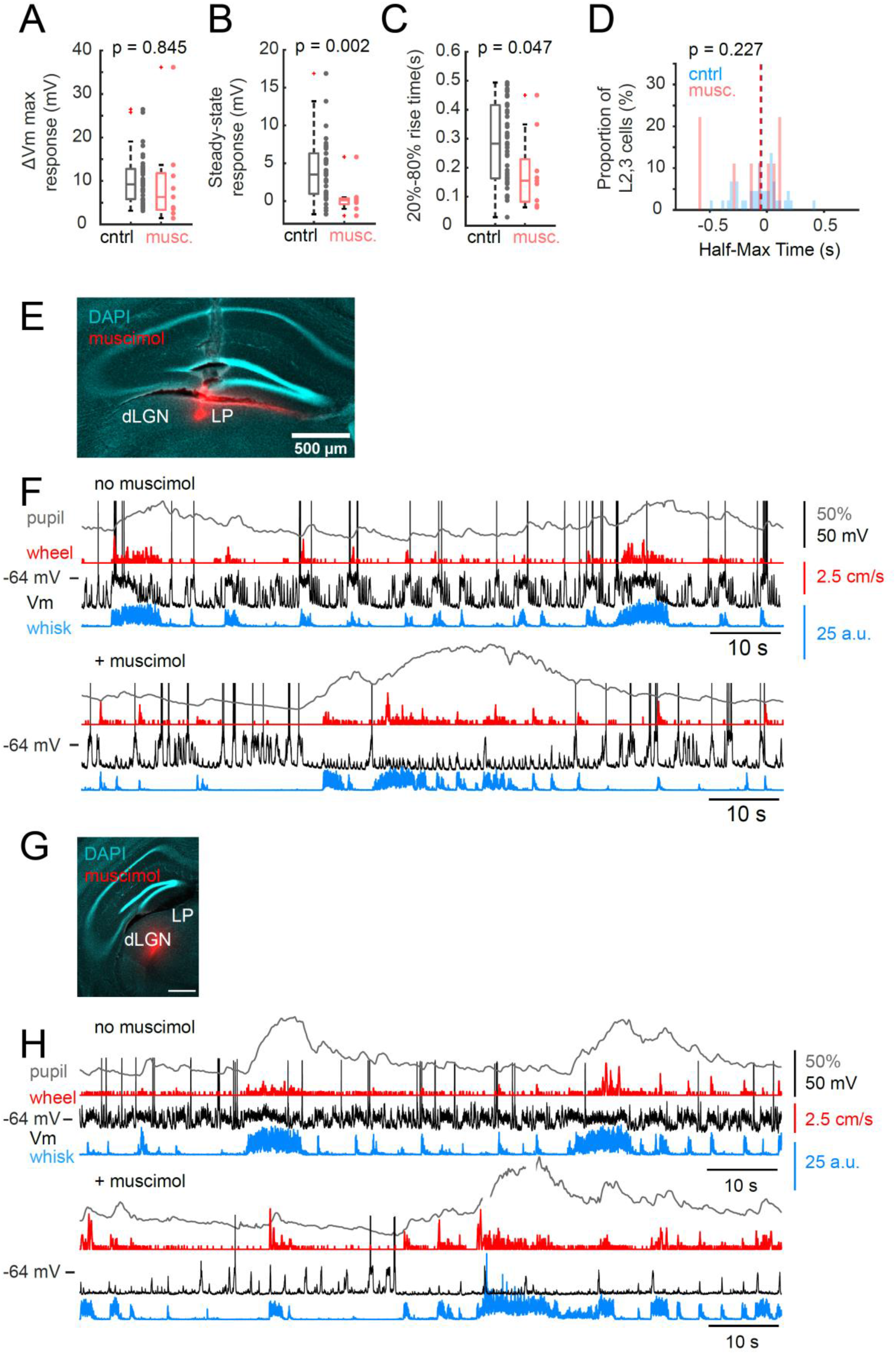
Effects of silencing of visual thalamus on movement/arousal related activity in V1. (A) The maximum response of the depolarization associated with locomotion in L2,3 V1 neurons is not significantly changed by the inactivation of the visual thalamus. (B) The steady-state response of the locomotion related response in L2,3 neurons is largely abolished by inactivation of the visual thalamus. (C) The 20%-80% rise time of these depolarizations is slightly changed but the (D) half max time remains unchanged by inactivation. (E) Example image of a muscimol injection that was centered around the LP nucleus. (F) The top trace shows a whole-cell recording of a L2,3 neuron before muscimol was injected into the visual thalamus (the same cell is also shown in Fig.S1A and S2A). The bottom trace depicts a whole-cell recording of a L2,3 neuron from the same animal as in the top trace and from the brain shown in (E) after muscimol was injected. (G) Example image of a muscimol injection that was centered around the dLGN nucleus. (H) The top trace shows a whole-cell recording of a L2,3 neuron before muscimol was injected into the visual thalamus. The bottom trace depicts a whole-cell recording of a L2,3 neuron from the same animal as in the top trace and from the brain shown in (G) after muscimol was injected.

